# From Correlation to Causation: Cell-Type-Specific Gene Regulatory Networks in Alzheimer’s Disease

**DOI:** 10.1101/2025.09.25.674923

**Authors:** Danni Liu, Zhongli Jiang, Hyunjin Kim, Anke M. Tukker, Ashish Dalvi, Junkai Xie, Yan Li, Chongli Yuan, Aaron B. Bowman, Dabao Zhang, Min Zhang

## Abstract

**INTRODUCTION:** Alzheimer’s disease (AD) involves complex regulatory disruptions across multiple brain cell types, yet a comprehensive understanding of the intracellular causal mechanisms remains unclear.

**METHODS:** We presented an integrative analysis framework using single-nucleus transcriptomic with matched subject-level genotype data from 272 human AD in the Religious Orders Study and the Rush Memory and Aging Project (ROSMAP) study, and constructed causality-based, cell-type-specific gene regulatory networks (GRNs).

**RESULTS:** Our method identifies regulatory genes from both transcription factors (TFs) and non-TFs, thereby capturing a complete and accurate causal regulatory map across different brain cell types. This work revealed both established and novel regulations, pathways, and cell-type-unique hub genes in AD. Beyond constructing transcriptome-wide GRNs, we quantitatively assessed hub genes and distinguished those with regulatory or responsive roles.

**DISCUSSION:** Our study provides a comprehensive mapping of cell-type-specific causal GRNs in AD, providing a powerful resource for dynamic pathway exploration, hypothesis generation, and functional interpretation.

## 1 Background

Alzheimer’s disease (AD), a complex neurodegenerative disorder characterized by progressive cognitive decline and memory loss, is the most common form of dementia and is predicted to affect 13.8 million Americans over the age of 65 by 2060.[1,2] Numerous genetic risk factors linked to AD have been revealed from previous genome-wide association analyses (GWAS), including the apolipoprotein (*APOE*) gene and the amyloid precursor protein (*APP*) gene. [1,3,4] Still, a major bottleneck for understanding AD pathogenesis lies in its biological complexity, involving intra- and inter-cellular interactions, neuronal loss, gliosis, and accumulation of pathological proteins, which are all modulated under complex gene regulatory networks (GRNs). [3,5] These precise regulatory mechanisms remain largely unknown, especially in terms of how they mediate expression changes in AD and how they vary between brain cell types. Therefore, understanding interactions between genes and transcription factors (TFs) within specific cell types is crucial for uncovering the cellular processes underlying neurodegeneration and AD progression, ultimately guiding the development of targeted treatments and preventive care for AD.

Classical methodologies to construct GRNs, such as the weighted gene co-expression network analysis (WGCNA) [6,7], rely on calculating correlations between genes using only transcriptomic data from bulk tissue. While these correlation-based networks are valuable in identifying co-expressed gene modules and potential biomarkers, they are limited in their ability to reveal the underlying causal interactions between genes, thereby hindering our understanding of the molecular mechanisms driving AD. [8] On the other hand, the advent of single-cell and single-nucleus sequencing technologies, along with the availability of large-scale genomic data, offers unprecedented opportunities to dissect causal interactions between genes at the resolution of cell type level by integrating transcriptomic and genomic information. Even though many studies have been carried out for constructing cell-type-specific GRNs using single-cell data, such as the widely used SCENIC [9] or master regulator analysis (MRA) in MARINa [10] and MOMA [11], they still primarily rely on correlation-based models and tend to focus narrowly on major TFs and their direct targets, potentially overlooking the regulations where non-TFs act as upstream regulators or the complex intergenic loops among TFs themselves. This gap leaves a large portion of unmapped or poorly understood regulations, thus bringing the need for a more comprehensive regulatory map that extends beyond the TF-centric assumptions and integrates causal inference to capture the complex gene regulations accurately.

In this study, we combined single-nucleus RNA sequencing (snRNA-seq) and whole-genome sequencing (WGS) data from the Religious Orders Study and the Rush Memory and Aging Project (ROSMAP) study to construct cell-type-specific causal GRNs for AD patients. We developed a comprehensive analysis pipeline to apply *SIGNET*[12] (<u>S</u>tatistical Inference on <u>G</u>ene Regulatory <u>Net</u>works), which employed a two-stage penalized least squares (2SPLS)[13] approach to conduct transcriptome-wide Mendelian Randomization (MR) with cell-type-specific expression quantitative trait loci (eQTLs) as instrumental variables. This framework allows us to move beyond the traditional correlation-based networks and TF-driven regulations to infer comprehensive, cell-type-specific, causal interactions between all genes in the AD brain. By identifying novel AD-associated hubs and key pathways as potential biomarkers, this study advances our understanding of the molecular mechanisms driving AD and holds enormous potential for developing targeted avenues of AD diagnosis and treatments. Moreover, the analytical pipeline can be broadly extended to other multi-omics data for the construction of cell-type-specific causal GRNs across diverse biological contexts.

## 2 Methods

### 2.1 Study Cohort

Individuals in this study were part of the multi-omics atlas of ROSMAP.[14] We extracted a subset of 424 individuals for whom both snRNA-seq and WGS data were available from dorsolateral prefrontal cortex (DLPFC) tissue.[15] A comprehensive quality control process was carried out to exclude low-quality and duplicated specimens, as well as individuals with distant ancestry as determined by a genotype Principal Component Analysis (PCA). (See details in **Supplementary Information S3**) Based on a neuropathological diagnosis using the total score derived from the Consortium to Establish a Registry for Alzheimer’s Disease (CERAD), 272 participants with pathological diagnoses of definite and probable AD were included for GRN construction.

### 2.2 Causal Gene Regulatory Network Construction Pipeline

To construct causality-based, cell-type-specific GRNs, we employed the *SIGNET* computational model to AD samples from the ROSMAP cohort. This approach leverages the power of genomic variants as instrumental variables to establish robust causal relationships between genes. The framework consists of four major steps, which will be detailed in subsequent sections: 1) preprocessing of snRNA-seq data for quality control, cell-type identification, and confounding adjustment; 2) preprocessing of WGS data for missing value imputation and genomic variant screening; 3) identification of cell-type-specific instrument variables (ctIV) through cis-eQTL mapping, stratified by minor allele frequencies (MAF) to ensure reliable analysis of rare alleles; and 4) construction of cell-type-specific causal GRNs using transcriptome-wide causal inference from snRNA-seq and WGS data. (**Figure 1A**)

**Figure 1.**
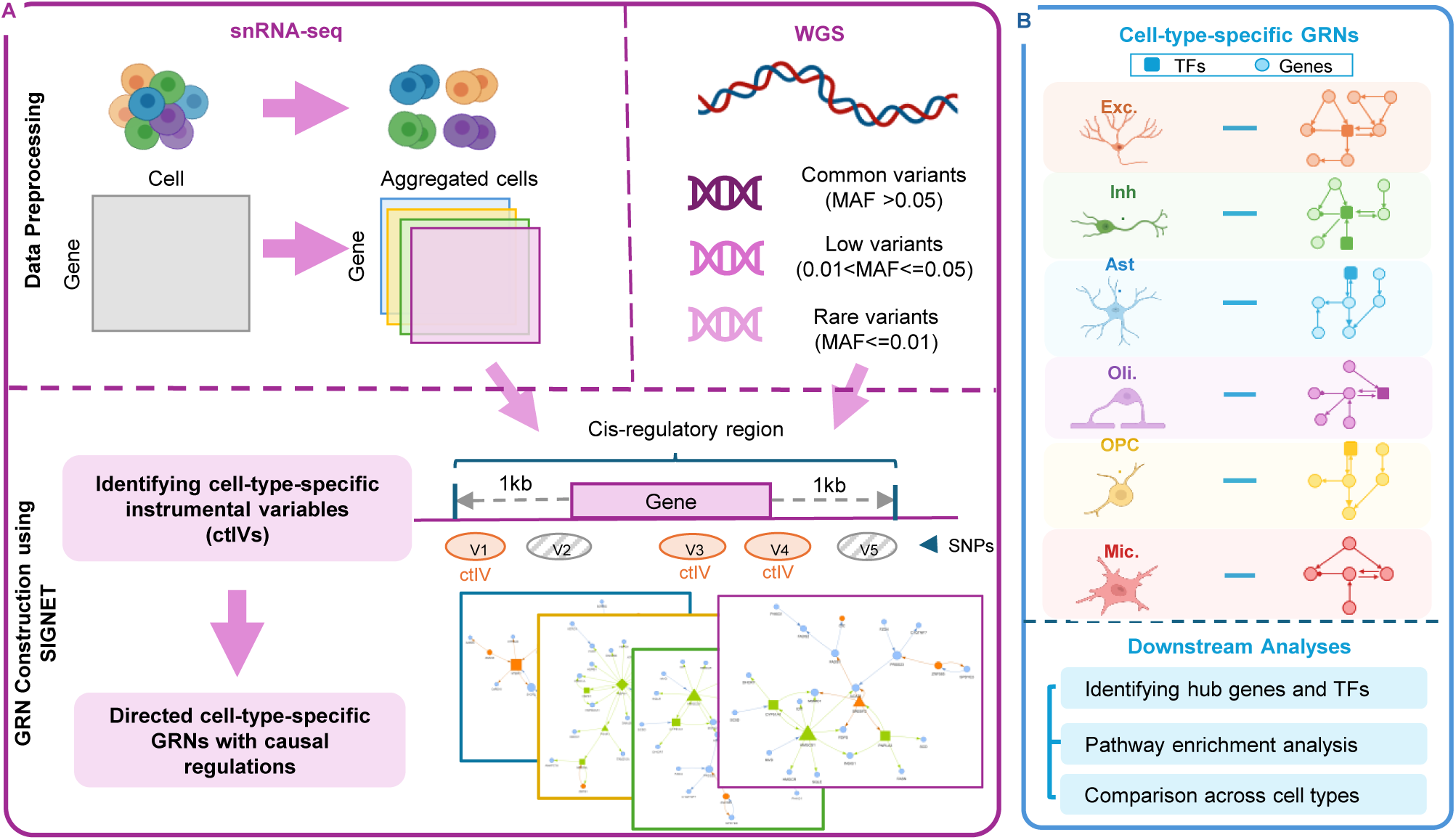
The pipeline of cell-type-specific causal GRN construction. (A) The workflow to construct cell-type-specific GRNs. It begins with the preprocessing of both snRNA-seq and WGS data, followed by the construction of GRNs using *SIGNET*. Detailed description of the workflow can be found in the **Methods section 2.2**. (B) Constructed cell-type-specific GRNs for each cell type. Downstream analyses were performed to decipher the complex regulatory relationships and identify hub genes of interest.

#### 2.2.1 Preprocessing of single-nucleus gene expression data

Single-nucleus RNA sequencing data underwent comprehensive preprocessing for quality control at the cell and gene levels, as well as cell type classification to characterize cell-population structures.[15] Quality control excluded cells with low sequencing depth, potential doublets, or dead cells, ensuring accurate cell classification and reliable inferred gene regulations. Gene expression profiles of filtered cells were normalized and scaled using *SCTransform* [16] in the *Seurat* package to prepare for cell-population classification.

#### 2.2.2 Cluster annotation and aggregation of single-nucleus data

Clusters of similar cells were detected using the Louvain algorithm on a k-nearest-neighbor (KNN) graph, constructed on a lower-dimension representation of top variable features obtained through PCA. [17] These clusters were then assigned to eight cell types (excitatory neurons (Exc.), inhibitory neurons (Inh.), astrocytes (Ast.), oligodendrocytes (Oli.), oligodendrocyte progenitor cells (OPC.), microglia (Mic.), endothelial cells, and pericytes) using a weighted elastic-net logistic regression, which acted as an automatic annotation classifier, with 24 donors from a previous DLPFC atlas as the reference. [15,18]. Endothelial cells and pericytes were excluded from further analysis due to an insufficient number of cells and a low identification rate within the cohort. For each identified cell type, the sparse single-nucleus gene expression data were aggregated by summing expression values across all cells of that type within each individual. Subsequent network construction was conducted independently for each cell type.

#### 2.2.3 Adjusting for covariates

To remove confounding effects and batch effects, we fitted a mixed-effects log-linear model using the aggregated individual-level counts of each gene. Confounding factors, including age, sex, postmortem interval (PMI), study cohort (ROS or MAP), and *APOE ε*4 genotypes, were included with fixed effects, while the batch effects of specimens were taken as random effects. The number of cell-type-specific nuclei in each individual was log-transformed and was included as an offset term to account for variations from cellular sampling depth. Finally, the working residuals were output for the downstream GRN construction.

#### 2.2.4 Genotype preprocessing and imputation

Whole genome sequencing data went through comprehensive quality control and imputation steps to prepare for GRN construction. First, we excluded multi-allelic or duplicating variants and kept SNPs with a missing rate under 0.1, minimum minor allele counts across all samples greater than 5, and not deviating from the Hardy-Weinberg equilibrium (HWE) test (p>1×10^-6^). Samples were then filtered to retain those with a missing rate below 0.2. All quality control procedures were carried out using *PLINK*[19] software. Afterwards, missing genotypes were imputed using the software *IMPUTE2* [20], with the 1000 Genome Phase 3 data as the reference panel. A second round of quality control was carried out on the imputed data, applying the same filtering criteria as in the first round with the additional requirement of an imputation quality control score greater than 0.5. Genotype data that passed both rounds of quality control were used for GRN construction.

#### 2.2.5 Construction of cell-type-specific GRNs with causal regulations

To construct causality-based, cell-type-specific GRNs, we extended the Mendelian randomization methodology as in our previous study of *SIGNET* [12] and applied it to the AD samples. This approach involves two stages: identifying cell-type-specific instrumental variables (ctIVs) through cis-eQTL mapping, and constructing cell-type-specific regulations of each gene through causal inference conducted using the identified ctIVs. The cis-eQTL mapping was performed with each gene’s cis-regulatory region defined as 1000 base pairs upstream and downstream of the gene’s genomic region, unless this overlapped with neighboring genes, in which case the overlapping region was divided between them. The variants within the region were categorized based on MAF into three levels: common variants (MAF > 0.05), low-frequency variants (0.01 < MAF ≤ 0.05), and rare variants (MAF ≤ 0.01). While common variants were directly taken for cis-eQTL mapping, the low-frequency and rare variants underwent an adaptive Sum (aSum) score test, where variants with similar effects were aggregated as combined ctIVs to enhance statistical power. To avoid redundancy, the top three most ctIVs with pairwise correlation under 0.9 were selected for each gene. At the causal inference stage, we implemented the two-stage penalized least squares (2SPLS) approach, first predicting the expression of potential regulators, i.e., genes with ctIVs, based on all potential regulators’ ctIVs using ridge regression, and then identifying significant regulations for each gene from potential regulators using an adaptive LASSO. [12] To ensure robustness in the constructed gene regulations, we reconstructed GRNs in 1,000 bootstrap datasets and retained only those regulations that appeared in at least 95% of the GRNs. If the final GRN was too extensive to be visualized, we further partitioned it into subnetworks based on modularity. [12] We focused all subsequent analyses on the top five largest subnetworks.

### 2.3 Identifying Hub Genes from GRNs

Characterizing each gene’s contribution in a large network is challenging. We are particularly interested in identifying hub genes with high connectivity, as they are likely to play central roles in regulating other genes or responding to regulatory signals. For each gene, we calculated the total number of regulations linked to it, referred to as degree, and further broke down this number into incoming and outgoing regulations, known as indegree and outdegree, respectively. We identified hub genes as those with a total degree exceeding the 95th percentile among all genes. To quantify both the GRN’s regulatory power on each gene and each gene’s influence on others, we defined two statistical measures, *R*^2^ and *C*^2^, measuring the extent to which a gene is determined by other genes and its influence on downstream genes, respectively (See details in **Supplementary Information S7**). [21] We calculated *C*^2^ as the summed proportions of variance a gene explains in all its target genes’ variance. A high *C*^2^ value indicates that a gene plays an important role in regulating its downstream genes in general. Conversely, we employed *R*^2^ to measure the proportion of a gene’s variance explained by all of its upstream genes. Among these parameters, both outdegree and *C*^2^ are positive exclusively for potential regulators, which can regulate genes that may themselves be potential regulators or not. We used a uniform threshold of 0.95 quantile for *R*^2^ and *C*^2^ for selecting the most important hub genes and characterizing their key roles. Based on these parameters, hub genes were classified into three categories: regulatory hubs, responsive hubs, or others. Genes with both *R*^2^and indegree exceeding 0.95 quantile of all genes are considered as “responsive hubs” that stand out from all regulated genes. Similarly, genes with both *C*^2^ and outdegree exceeding 0.95 quantile of all genes are classified as “regulatory hubs” that stand out from all gene regulators.

### 2.4 Pathway Enrichment Analysis and IPA

To understand the biological processes and pathways revealed in the constructed GRNs and the cell-type-specific activities, we performed pathway enrichment analysis for each cell type on its complete network, as well as its five largest subnetworks. Enrichment tests were carried out using the R package *clusterProfiler* [22] with Fisher’s exact test. Benjamini-Hochberg correction was applied to calculate the adjusted p-values for multiple tests, and significantly enriched pathways were selected with adjusted p-values under 0.05. We selected the top 20 enriched pathways by also considering the gene ratio, defined as the proportion of genes in our network annotated to a specific pathway.

### 2.5 IPA and UCI ADRC Multiomics Cohort

For additional validation and cross-checking with existing literature, the lists of genes identified in our network were submitted to QIAGEN Ingenuity Pathway Analysis (IPA) [23] (QIAGEN Inc., https://digitalinsights.qiagen.com/IPA) software to look for canonical pathways or regulations supported by existing literature.

We further evaluated the identified regulations in our networks using an independent cohort from the University of California Irvine (UCI) Institute for Memory Impairments and Neurological Disorders (MIND)’s Alzheimer’s Disease Research Center (ADRC) Multiomics Study. [24] This cohort includes prefrontal cortex samples from 11 individuals with late-stage AD, and cells were classified into the same six major cell types as in the ROSMAP study. Spearman’s correlation coefficients were calculated to validate the identified regulations between 20 AD causal genes reported by the Alzheimer’s Disease Sequencing Project (ADSP) and their directly linked genes.

## 3. Results

### 3.1 Study Cohort and Analysis Pipeline

In this study, we utilized snRNA-seq data alongside WGS data collected from frozen DLPFC tissue samples in ROSMAP, a pivotal cohort study designed to advance the understanding of AD and other chronic neurological conditions associated with aging. After quality control, 272 individuals who were pathologically diagnosed with AD using the CERAD score were selected for network construction [25,26] **(Table 1).** This cohort is predominantly female (71.69%), with balanced distributions across both ROS and MAP study projects. Over half of the cohort (51.84%) were over 90 years old, with the remainder aged above 71.31 years and averaging

**Table 1.** Demographic and Clinical Characteristics of Study Participants. Summary statistics of demographics for 272 AD participants in the ROSMAP cohort that is used in this study. All participants have both single-nucleus RNA-seq and genotype data available and passed quality control. All participants are white and non-Hispanic.

85.11 years. All samples in this cohort are white and non-Hispanic. The APOE *ε*4 alleles showed a high prevalence among the AD participants, with 31.62% samples carrying one *ε*4 allele and 1.47% samples carrying two *ε*4 alleles. Conversely, the APOE *ε*2 allele was absent in the majority of the cohort, with 85.29% of the participants not possessing any APOE *ε*2 allele, and only 14.7% carrying one allele.

To construct cell-type-specific GRNs, we developed a comprehensive analysis pipeline that incorporated both snRNA-seq and WGS data within a Mendelian randomization framework to infer causal gene regulations for each cell population. (**Figure 1**) After quality control of snRNA-seq data, we obtained 947,268 nuclei in DLPFC samples across 272 AD participants, with each individual having an average of 3,483 nuclei. These nuclei were classified into six major brain cell types, including excitatory neurons (Exc.; 375,092 nuclei), inhibitory neurons (Inh.;148,224 nuclei), astrocytes (Ast.;131,008 nuclei), microglia (Mic; 49,263 nuclei), oligodendrocytes (Oli.; 202,714 nuclei), and oligodendrocyte progenitor cells (OPC.; 34,936 nuclei). (**Figure 2A**) The cell counts in different cell populations are distributed evenly across subjects, with excitatory neurons being the largest cell population in all individuals.

**Figure 2.**
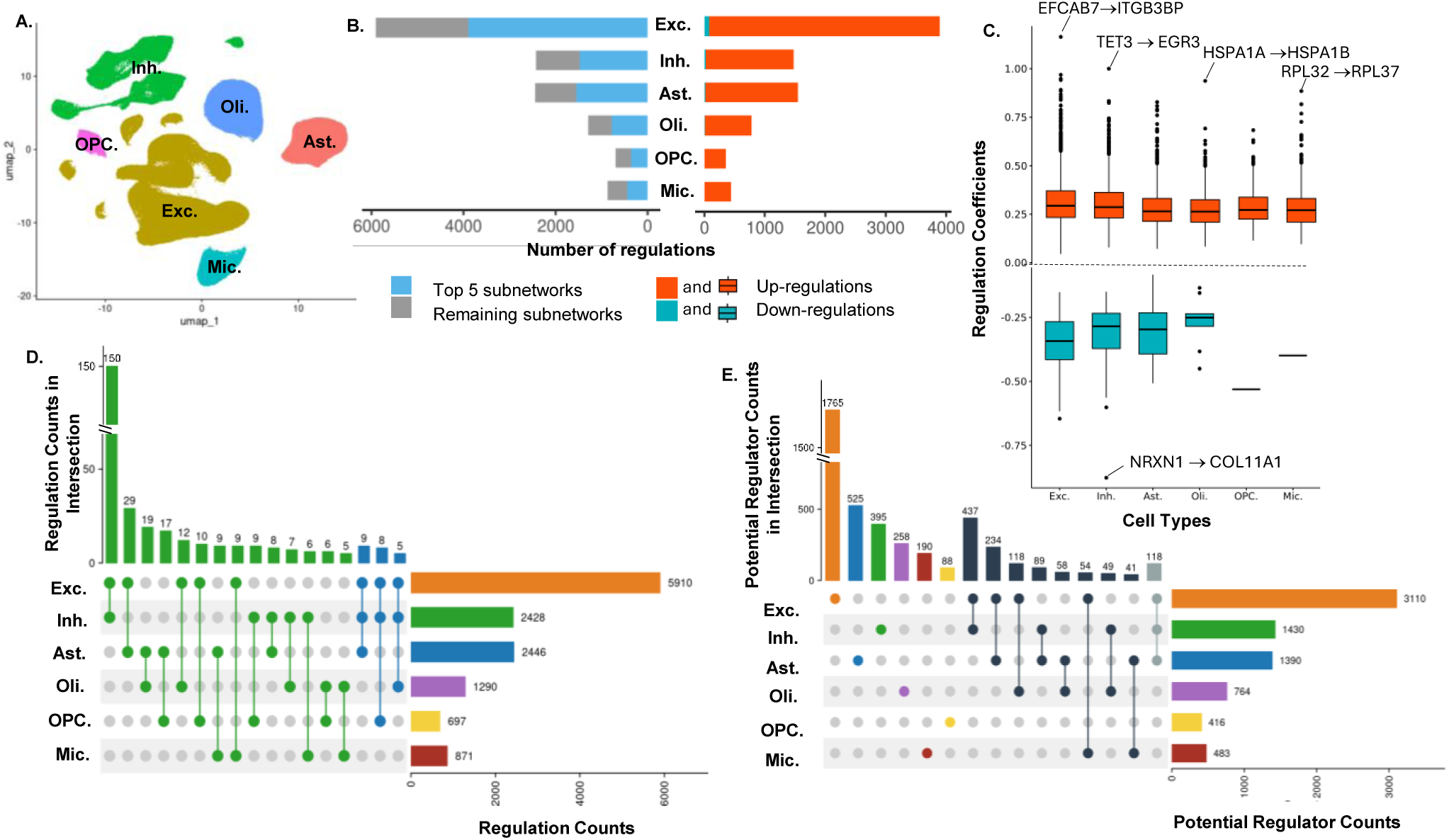
Global characteristics of cell-type-specific GRNs in AD. (A) UMAP plot of six cell populations from snRNA-seq data of 272 AD patients. (B) Summary of regulation counts in GRNs across different cell types. The length of bars in both right and left panels indicates the regulation counts in the cell types. In the right panel, the regulation counts from the top five largest modules were highlighted in blue (see exact counts in **Supplementary Information Table S1**). In the left panels, the regulations were categorized into down- and up-regulations. (C) Boxplots of effect sizes for up- and down-regulations across cell types. The effect size indicates the per-unit change in the standardized gene expression. The top five regulations with the largest absolute effect sizes were labeled. (D) Regulations shared across cell types. The top panel shows counts of shared regulations across pairs (green) or triplets of cell types (blue) in the left panel. Cell-type combinations not shown have fewer than five shared regulations. The right panel shows per-cell-type regulation counts. (E) Potential regulators across cell types. The left panel shows per-cell-type counts of potential regulators (in the first six bars) and counts of potential regulators shared by different pairs of cell types (in the other eight bars). Cell-type combinations not shown have fewer than 30 shared potential regulators. The right panel shows per-cell-type counts of potential regulators.

### 3.2 Global Characteristics of Cell-Type-Specific GRNs in AD cell types

The constructed GRNs reveal variations in network complexity across neurons and glial cells. Excitatory neurons exhibited the most extensive GRN with 5,910 regulations, followed by inhibitory neurons with 2,428 regulations. Among glial cells, astrocytes had the largest network with 2,446 regulations, while microglia and OPCs displayed the sparsest network among all cell types. (**Figure 2D, Supplementary Information Table S1**) We classified genes with ctIVs as potential regulators, whose regulatory effects can be inferred by multiple Mendelian randomization. While potential regulators comprise 52-62% of the genes in the GRNs, 42-49% of regulations occur between potential regulators themselves. A closer investigation of the regulatory effects revealed that positive regulations dominated the identified regulatory networks across all cell types. (**Figure 2B**) Particularly, excitatory neurons exhibited the largest variance in the regulatory effects in both directions, with the highest number of 103 down-regulations and 5,807 up-regulations, among which 441 regulations have coefficients greater than 0.5. Despite microglia and oligodendrocytes having smaller overall network sizes compared to the neurons, the strongest regulations (*HSPA1A* to *HSPA1B* in oligodendrocytes; *RPL32* to *RPL37* in Mic.) have effect sizes of 0.94 and 0.89, respectively, ranking among the top five regulations with the largest absolute effect sizes across all cell types. (**Figure 2C**) The strongest up-regulation effect, with an effect size of 1.16, was observed in excitatory neurons from gene *EFCAB7* to *ITGB3BP*, followed by inhibitory neurons with strong regulations in both directions: up-regulation of *TET3* to E*GR3* (effect size = 1) and down-regulation of *NRXN1* to *COL11A1* (effect size = 0.88).

We performed a comparison of GRNs constructed across cell types to reveal shared and cell-type-specific regulatory patterns. A comparison of regulations showed the highest number of 150 shared regulations between the two types of neuronal cells, followed by 29 regulations between excitatory neurons and astrocytes. (**Figure 2D**) The same pattern was also observed in the comparison of potential regulators across cell types, with 437 potential regulators shared between two neuron types, followed by 234 potential regulators between excitatory neurons and astrocytes. This comparison also revealed unique potential regulators in each cell type. Excitatory neurons possessed the largest number of 1,765 exclusive potential regulators, comprising 56.75% of all potential regulators in their network, while the composition is much lower in other cell types, ranging from 21.15% to 39.33%.

### 3.3 Identification of GRN Hubs Across Cell Types

Due to the extensive size of the full networks, we further separated the networks into subnetworks to allow for easier visualization. Counting regulations in each cell type, the top five subnetworks accounted for 50.93-65.89% of the complete networks, with the excitatory neurons showing the largest proportion. (**Supplementary Information Table S1**) We focused our subsequent analyses on these subnetworks, since they captured the most complex and concentrated network structures. Across the six cell types, we investigated the hub genes, some of which are further categorized into regulatory and responsive hubs. (**Figure 3A**)

**Figure 3.**
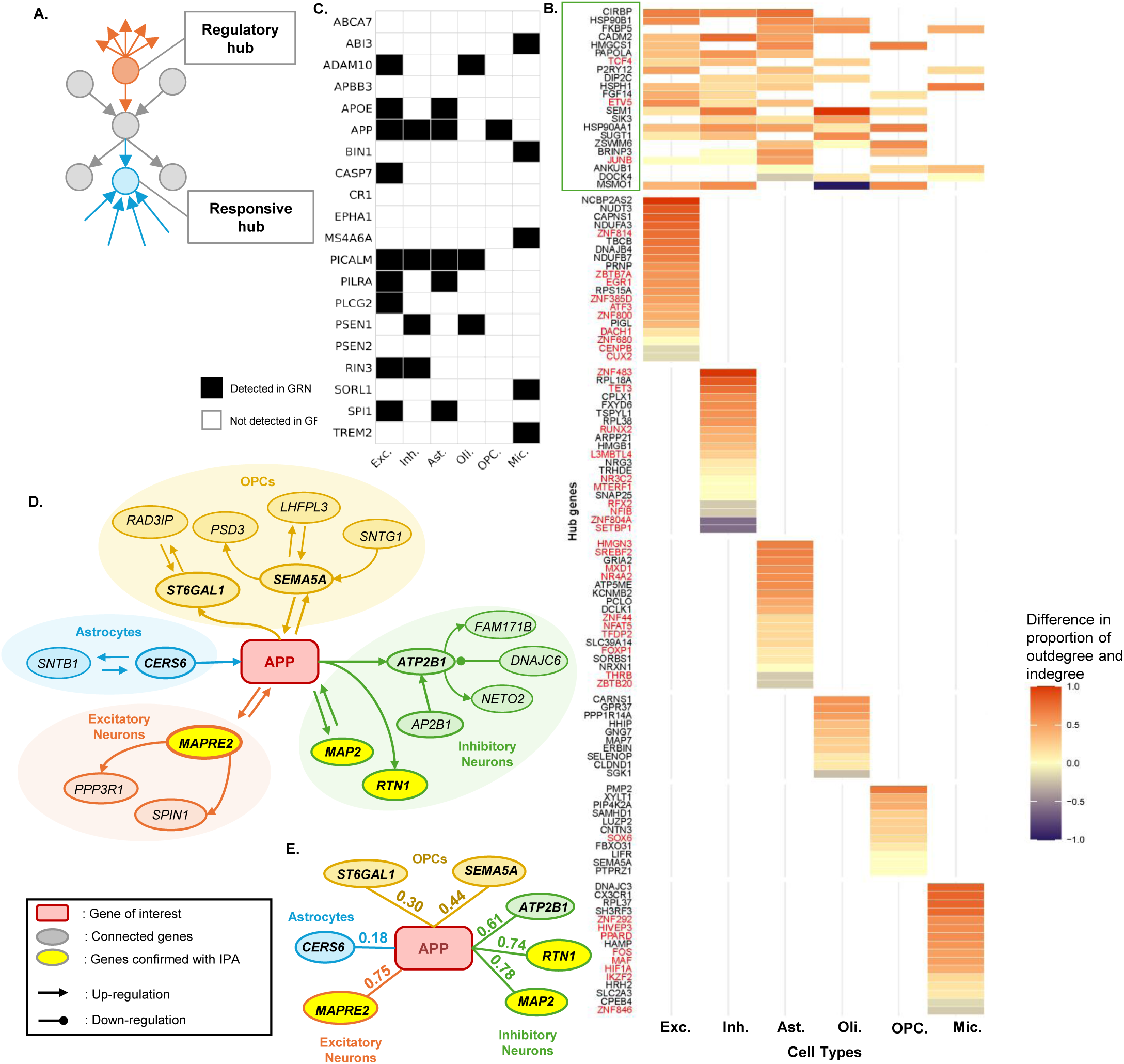
Hub genes and AD-risk causal genes across cell types. (A) Illustration of types of hub genes. Regulatory hubs and responsive hubs were further selected to have their outdegrees and indegrees ranking in the top 5%, respectively. (B) Distribution of hub genes across cell types, including the top 10 cell-type-unique hub genes, the top 20 cell-type-unique TF genes, and hub genes shared by more than two cell types (frame green). Color represents the difference between outdegree and indegree proportions for each gene. TF gene names are highlighted in red on the Y-axis. Complete data of this figure can be provided in **Supplementary Information Table S3**. (C) Appearance of 20 AD-relevant genes, reported by ADSP [33], in our constructed GRNs. *APP* and *PICALM* appear most frequently, each in four cell types. (D) *APP-involved* regulations across cell types, with genes confirmed to have connections in *QIAGEN IPA* highlighted in yellow. (E) Correlation among genes directly linked to *APP* using snRNA-seq data from an independent cohort.

We observed distinct sets of hub genes as well as different regulatory connectivity and directions across cell types. Excitatory neurons contained the largest set with 427 hub genes, followed by 178 hub genes in astrocytes. Across cell types, 62.06-81.13% of hub genes are regulatory hubs, indicating that the majority of the hub genes serve as key regulators of downstream gene activities. TF genes constitute 10.23% of hub genes on average, with a minimum of 5.68% of hubs being TFs in oligodendrocytes and a maximum of 15.25% in microglial cells. Specifically, a large proportion (48.65 −75%) of the TF hubs were identified as regulatory hubs, reflecting their consistently regulatory role across all cell types. On the contrary, responsive hubs, which primarily react to upstream regulatory signals, were comparatively rare, and those that were also TFs appeared in only several cell types. In fact, inhibitory neurons and astrocytes have the largest set, each with only three TFs as responsive hubs (inhibitory neurons: *ZNF804A, NFIB, SETBP1;* astrocytes: *THRB, ZBTB20, TRPS1*), while none of the TFs were responsive hubs in excitatory neurons. (**Supplementary Information Table S2**)

There were no hub genes shared across all cell types nor among all glial cells, while a total of 110 hub genes, including 14 TFs (top three TFs: *EVT5, JUNB, TCF4*), were shared among multiple cell types. (**Figure 3B, Supplementary Information Table S3**) Specifically, the two types of neuronal cells share 43 hub genes, with 10 genes appearing among the top 20 hub genes in both cell types, i.e., RPS27A, *HSP90AA1, MSMO1, PAPOLA, CIRBP, JUNB, ETV5, HSPH1, DPH6, and GNAS*. (**Figure 3B**). Among them, genes *RPS27A, HSP90AA1, MSMO1, PAPOLA,* and *CIRBP* are the top five regulatory hubs in both cell types. *JUNB*, *ETV5,* and *TCF4* are the only TF genes in this list, with *JUNB* having balanced indegree and outdegree in both cell types. At the same time, *ETV5* is a top regulatory hub only in excitatory neurons, with outdegree eight versus indegree two. (**Supplementary Information Table S3**) **Figure 3B** also revealed sets of cell-type-unique hubs and TF genes that were potential cell-function-specific AD markers or AD-driving genes. For example, microglia contained 16 cell-type-unique hub genes, with gene *DNAJC3* having 10 among 11 regulations as outgoing regulations, followed by *CX3CR1 and RPL37* with outdegree nine among 10 regulations. Eight of these 16 hubs are cell-type-unique TF genes, with the top two TF genes, *FOS* and *MAF,* showing much larger outdegree than indegree, while *ZNF846* showed higher indegree than outdegree. (Supplementary Information Table S3)

### 3.4 Prevalence of AD Causal Genes and Their Regulations

We further examined our constructed GRNs for the 20 AD-risk genes reported by the ADSP, including *ABCA7*, *ABI3*, *ADAM10*, *APBB3*, *APOE*, *APP*, *BIN1*, *CASP7*, *CR1*, *EPHA1*, *MS4A6A*, *PICALM*, *PILRA*, *PLCG2*, *PSEN1*, *PSEN2*, *RIN3*, *SORL1*, *TREM2,* and transcription factor *SPI1* [27–33]. Excitatory neurons contained the most, with nine AD-risk genes, followed by astrocytes and microglia, each with five genes. Genes *APP* and *PICALM* were identified in four cell types. (**Figure 3C**) In fact, *APP*, a crucial gene in AD due to its potential impact on *Aβ* peptide, was identified as a key modulator by IPA, while it is a strong regulatory gene in our constructed networks by explaining 73.01% of RTN1’s variation and 43.40% of MAP2’s variation in inhibitory neurons. (**Supplementary Information Table S2**) We examined *APP*’s regulations and its depth-2 neighbors across cell types in neurons, astrocytes, and OPCs, and cross-checked them with the IPA database. (**Figure 3D**) In excitatory neurons, our network revealed bidirectional causal regulation between the genes *APP* and *MAPRE2*, with each explaining 45.27% and 47.54% of the variation in the other. Whereas IPA indicated only an indirect connection between them, mediated by *SNCA,* which in our network acts only as a target gene, with 84.43% of the variation explained by upstream genes. (**Supplementary Information Figure S1A, Table S2**) Similarly, our GRN for inhibitory neurons revealed a bidirectional regulation between *APP* and *MAP2,* as well as a causal regulation of *RTN1* by *APP*. IPA results supported the interaction between *APP* and *MAP2* through protein-protein interactions mediated by *GRB2*, whereas the connections between *APP* and *RTN1* appear more complex, involving multiple intermediate proteins. (**Supplementary Information Figure S1B**)

An independent cohort containing 11 late-stage AD DLPFC samples from the UCI ADRC Multi-omics Study [24] was used to validate the regulations in our networks involving genes directly connected to the gene *APP*. The single-nucleus RNA-seq dataset contained 36,114 genes and 61,472 cells after quality control, which were further classified into the same cell types as in the ROSMAP study. (**Supplementary Information Figure S2**) This independent snRNA-seq data showed strong correlations for *APP*-related regulations in excitatory and inhibitory neurons, with the highest correlation of 0.78 observed between *APP* and *MAP2* in inhibitory neurons. On the contrary, the correlations were weaker in astrocytes and oligodendrocytes, with astrocytes showing a correlation of 0.18 between *APP* and *CERS6,* and oligodendrocytes showing correlations of 0.30 and 0.44 between *APP* and gene *ST6GAL1* and *SEMA5A*, respectively. Notably, the three genes (*MAPRE2* in Exc. and *MAP2* and *RTN1* in Inh.), whose regulations were confirmed by IPA, showed the strongest correlations above 0.7 in this independent cohort. (**Figure 3E**)

### 3.5 The Excitatory-Neuron-Specific GRN

Since the excitatory neuron GRN exhibited the most extensive regulatory relationships and AD-relevant pathway activities, we further examined it to better understand the cellular activities of this cell type. The largest subnetwork of excitatory neurons contained 3,662 regulations and involved 427 hub genes (11.66%), comprising 265 regulatory hubs and 21 responsive hubs. (**Supplementary Information Table S2**) Among all hub genes, 37 (8.67%) are TF hubs, among which 18 (48.65%) are regulatory hubs, with the top five being *ETV5, EGR1, NR4A1, ZBTB7A, and ZNF385D,* while none of these TFs were responsive hubs. Specifically, three of the top five TF hubs, *EGR1, ZBTB7A,* and *ZNF385D,* were also excitatory-neuron-unique hubs. The top five non-TF excitatory-neuron-unique hubs are *NDUFA3, PIGL, TBCB, PRNP, and NUDT3*, all being regulatory hubs. (**Supplementary Information Table S3**) This indicates the distinctive role of hub genes in driving the activities of other genes rather than being targets of others.

A closer investigation of excitatory neurons’ hub genes and TFs’ physical locations revealed distinct genomic hotspots, which contain numerous hub genes that regulate important cell functions. (**Figure 4A**) Gene *RPS27A* is the only and largest hub gene identified on chromosome 2 with the densest regulatory activities that outperform other hub genes. The most notable genomic hotspot was observed on chromosome 19 with 20 hub genes, led by gene *NDUFA3,* which has the second largest degree of 20. This chromosome has three TF hubs *JUNB, ZBTB7A,* and *ZNF814*, where *JUNB* is the largest TF hub while *ZBTB7A* and *ZNF814* are among the excitatory neuron-specific hubs. The second largest TF hub *ETV5* was located on chromosome 4, and both chromosomes 14 and 16 contain multiple hub genes with moderate to high degrees above the threshold. Besides, the largest responsive hub gene *UBR5* was found on chromosome 8, with six of the nine regulations as incoming regulations.

**Figure 4.**
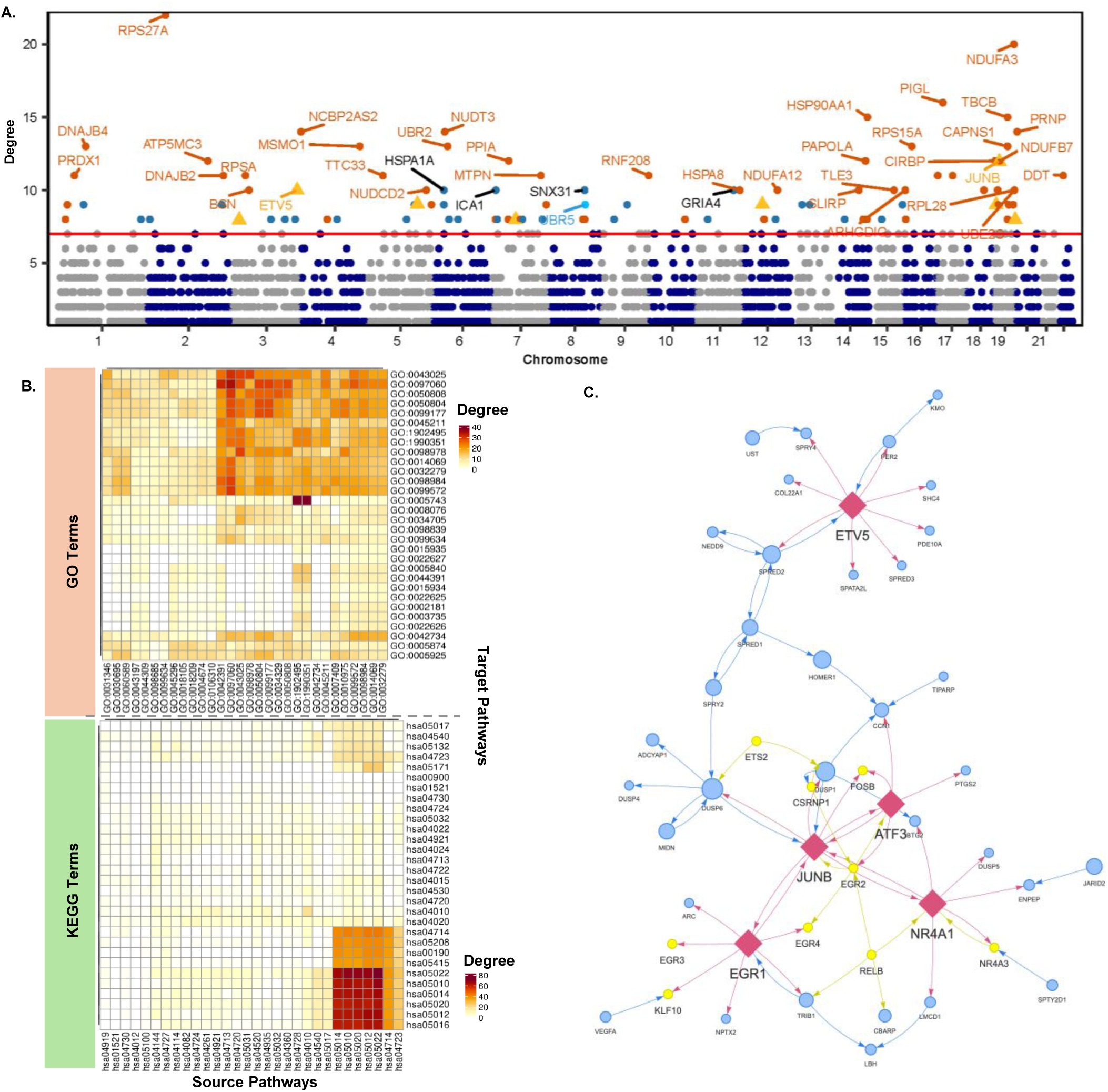
A closer investigation of the excitatory-neuron-specific GRN. (A) A Manhattan plot showing the physical distribution of all genes in the constructed excitatory-neuron-specific GRN. Genes in different chromosomes were colored in alternating colors. The Y-axis indicates the total degree of each gene. The red horizontal line is the 0.95 quantile degree of all genes in the GRN, serving as a cutoff. Genes with a degree higher than the 0.95 quantile cutoff were highlighted in the figure. Non-hub genes were highlighted in blue, regulatory hubs with the top 0.95 quantile of outdegree and *C*^2^ were highlighted in orange, and responsive hubs with top 0.95 quantile of indegree and *R*^2^ were highlighted in sky blue. TF hubs were highlighted in yellow triangles. Only hubs greater than the 0.99 quantile were labeled on this figure to avoid too many overlaps. (B) Heatmaps showing the pathway enrichment signals from the direction of upstream regulators to their targets for both GO and KEGG pathways. Only the top 30 significantly enriched pathways from pathway analysis were included in the heatmap, with rows as pathways enriched with source genes and columns as pathways enriched with target genes. The blocks in the heatmap were colored based on the number of regulations between the corresponding pair of pathways. Darker colors indicate more regulations. Corresponding pathway names of the IDs can be found in **Supplementary Information Table S4**. (C) A zoom-in GRN figure for selected TF hubs (JUNB, ETV5, EGR1, ATF3, and NR4A1) and their two nearest neighbors. The TFs were highlighted in pink diamond shape, with non-hub TFs in yellow circles and the rest in blue circles. The size of the circles is determined by the gene’s degree.

Pathway enrichment heatmap of excitatory neurons revealed directional signals from upstream to downstream genes in concentrated GO and KEGG pathways. (**Figure 4B**) The most prominent regulatory activities were observed from the transporter complex to the mitochondrial inner membrane in GO pathways with 39 regulations. It also highlighted intensive interconnections among pathways, including synaptic organization (synaptic membrane, neuron-to-neuron synapses, postsynaptic density, etc.), neuronal cell body, and glutamatergic synapse, with each pair of pathways having 25-35 regulations. Notably, the KEGG pathway analysis reveals a strong enrichment in pathways of neurodegenerative diseases, especially AD and Parkinson’s disease, which supports the finding that excitatory neurons are highly involved in AD. (**Figure 4B, Supplementary Information Figure S3**)

We further investigated the regulatory networks surrounding the TF hubs and their two nearest neighbors. A tightly interconnected regulatory circuit was observed among five TF hubs, *JUNB, ETV5, EGR1, ATF3,* and *NR4A1*, with four TF hubs (*JUNB, EGR1, ATF3,* and *NR4A1*) directly interacting with each other while they connect with *ETV5* distantly through several intermediate genes. (**Figure 4C**) These interconnected TF hubs also extensively interact with many secondary TFs and simultaneously coordinate the rest of the genes. Separate pathway analysis of these TF hub genes and neighbors revealed activities from negative regulation or phosphorylation. (**Supplementary Information Figure S4, Table S5)**

## 4. Discussion

In this study, we constructed cell-type-specific causal GRNs to investigate the cellular regulatory activities across different brain cell populations by integrating snRNA-seq and WGS data from DLPFC samples of 272 AD patients in the ROSMAP cohort. By utilizing MAF-stratified cis-regulatory variants as cell-type-unique instrumental variables within a transcriptome-wide Mendelian randomization model, we moved beyond the traditional correlation-based methods. We established causal regulations with *SIGNET*, which enriches our knowledge of the cellular regulatory mechanisms for AD. The underlying algorithm is computationally efficient with parallel computing, enabling the use of bootstrapping to identify regulations that are consistently constructed across bootstrapped GRNs, thereby significantly reducing false regulations.

Our comprehensive analysis of cell-type-specific causal GRNs revealed excitatory neurons as exhibiting the most extensive regulatory network and greatest diversity in regulatory effects, with regulatory hubs predominantly represented among hub genes. The pathway analysis results also confirmed their central role in synaptic dysfunctions associated with AD, which were mainly driven by interruption from upstream hub genes.[34–38] Besides, linking gene functions to shared and cell-type-unique hub genes in excitatory neurons facilitates our understanding of cellular regulatory activities related to AD pathogenesis. In fact, *RPS27A* was identified as the largest regulatory hub gene unique to excitatory neurons, with the highest values in both degree and *C*^2^, which aligns with a recent study of protein-protein interaction networks of AD patients that identified *RPS27A* as one of the four critical hub genes.[39] This gene encodes the ribosomal protein S27A, which serves as a structural constituent of the 40S ribosomal subunit and has established roles in activating inflammatory activities in microglial cells, cell-cycle regulation, and proteasome-mediated protein degradation from past studies of neurodegenerative diseases and mild cognitive impairment progression, which provide compelling evidence that *RPS27A* functions as a regulatory hub. [40–42] Similarly, our network identified a mitochondrial complex surrounding gene *NDUFA3* on chromosome 19, which supports the role of mitochondrial dysfunction in AD from past transcriptomics studies.[43–45] Further investigation on other cell-type-unique hub genes could provide insights into potential biomarkers with specialized AD regulatory mechanisms.

Our analysis also extends beyond master regulator analysis (MRA) used in MARINa/VIPER [10] or other widely used GRN methods (SCENIC [9], GENIE3 [46], etc.), which prioritize TFs as hub genes and primarily emphasize their role as regulators. Our results suggest that TFs not only act as regulators, but also act as targets or intermediate genes that interconnect with other TFs or key regulators to modulate dynamic cellular activities. This complex interaction is supported by the regulatory circuit shown around TF genes *JUNB, ETV5, EGR1, ATF3,* and *NR4A1* in our network. (**Figure 4C**) Besides, our results also reveal that regulatory activities from non-TF genes are more extensive than those in TF genes. These non-TFs, such as *RPS27A* and *NDUFA3*, are major regulatory hubs regulating key cellular activities and maintaining cell functions; therefore, they should not be overlooked in regulatory network studies. Furthermore, our causal network analysis clarifies the ambiguous protein-protein interactions or co-expression correlations inferred from traditional network and pathway analysis tools like IPA [23], and ultimately advances our knowledge of shared and unique regulatory patterns in AD progression across different brain cell populations.

Moving forward, we will dive deeper into the current results to investigate networks involved in AD-specific pathologies across different cell types. We plan to perform differential gene regulatory analysis between AD and healthy samples in these pathways to identify AD-specific regulatory patterns. This comparison will allow us to distinguish the regulation changes involved in neurodegeneration from normal cell activities during aging. Additionally, we will extend this framework to an integrative analysis of snRNA-seq and single-nucleus Assay for Transposase-Accessible Chromatin sequencing (snATAC-seq), incorporating chromatin accessibility information. Studying the dynamics of regulatory networks during disease progression from Mild Cognitive Impairment (MCI) to late-onset AD will be particularly valuable in identifying biomarkers for early detection and interventional treatments.

## Acknowledgements

The results published here are in whole or in part based on data obtained from the AD Knowledge Portal Community Data Contribution Program. This research was partially supported by NIH grants R01AG080917, R01AG080917-02S1, R01GM131491, and R01GM131491-02S2, NCI grant P30CA062203, and UCI Anti-Cancer Challenge funds from the UC Irvine Comprehensive Cancer Center. The authors acknowledge the support of the Chao Family Comprehensive Cancer Center Biostatistics Shared Resource, which is supported by the National Cancer Institute of the National Institutes of Health under award number P30CA062203. The content is solely the responsibility of the authors and does not necessarily represent the official views of the National Institutes of Health or the Chao Family Comprehensive Cancer Center.

## Consent statement

All data used in this study were requested and approved from the AD Knowledge Portal for the ROSMAP study (syn3219045), with informed consent obtained from all human subjects.

## Availability of data and materials

Raw and processed single-nucleus RNA sequencing data and whole-genome sequencing data for samples used for this study can be accessed from the ROSMAP consortium on Synapse (https://www.synapse.org/#!Synapse:syn31512863). Codes will be made available after publication.

## Abbreviations

2SPLS: Two-stage penalized least squares
Aβ: Amyloid β
AD: Alzheimer’s disease
ADSP: Alzheimer’s Disease Sequencing Project
*APOE*: The apolipoprotein gene
*APP*: The amyloid precursor protein
aSum: An Adaptive Sum Score test
CERAD: Consortium to Establish a Registry for Alzheimer’s Disease
ctIV: Cell-type-specific instrumental variable
DLPFC: Dorsolateral prefrontal cortex
eQTL: Expression quantitative trait locus
FDR: False discovery rate
GCV: Generalized cross-validation
GLMM: Generalized linear mixed model
GO: Gene Ontology
GRN: Gene regulatory network
GWAS: Genome-wide association study
HWE: Hardy-Weinberg equilibrium
IPA: Ingenuity Pathway Analysis
KEGG: Kyoto Encyclopedia of Genes and Genomes
LASSO: Least Absolute Shrinkage and Selection Operator
MAF: Minor allele frequency
MAP: Rush Memory and Aging Project
MCI: Mild Cognitive Impairment
MIND: Institute for Memory Impairments and Neurological Disorders
MMSE: Mini-Mental State Examination
MRA: Master regulator analysis
PMI: Postmortem interval
PCA: Principal Component Analysis
ROS: Religious Orders Study
MAP: Rush Memory and Aging Project
ROSMAP: Religious Orders Study and Rush Memory and Aging Project
SIGNET: <u>S</u>tatistical Inference on <u>G</u>ene Regulatory <u>Net</u>works
snRNA-seq: Single-nucleus RNA sequencing
snATAC-seq: Single-nucleus Assay for Transposase-Accessible Chromatin sequencing
TF: Transcription factor
UMI: Unique Molecular Identifier
WGS: Whole-genome sequencing

## Abbreviations for cell types

Ast.: Astrocytes
Exc.: Excitatory neurons
Inh.: Inhibitory neurons
Mic.: Microglia
Oli.: Oligodendrocytes
OPC.: Oligodendrocyte progenitor cells

