## Supplementary Information for "From Correlation to Causation: Cell-Type-Specific Gene Regulatory Networks in Alzheimer’s Disease"

### S1. Study Cohort and Cell Populations

All patients used in this study were from a multi-omics atlas of the Religious Orders Study (ROS) and the Rush Memory and Aging Project (MAP), which is a pivotal cohort study designed to advance the understanding of Alzheimer's disease (AD) and other chronic conditions associated with aging. This study engaged more than 3,000 participants aged 65 and above from Catholic communities across the US and other individuals from retirement communities in northeastern Illinois.[1] Systematic profiling of multiple omics data was collected from the dorsolateral prefrontal cortex (DLPFC) of autopsied individuals, including whole-genome sequencing, RNA sequencing, DNA methylation, metabolomics, proteomics, and imaging data.[2] In this study, we selected 424 participants who had both single-nucleus RNA sequencing (snRNA-seq) data alongside whole genome sequencing (WGS) data available from frozen DLPFC tissue samples.[3] After quality control, 272 participants with pathological diagnoses of definite and probable AD were kept for the downstream analyses.

### S2. Data Quality Control

We performed comprehensive quality control at the individual level and the sample specimen level to ensure the highest data quality. The first round of quality control was performed on an individual level, where we explored the demographic data and excluded any individual with outlying characteristics that are different from the cohort. Genotype data of all participants were examined using Principal Component Analysis (PCA) to identify individuals with distant ancestry in addition to self-reported ethnicity information. As a result, six people were excluded from further analyses in this first round of quality control, including one individual of Hispanic origin using self-reported ethnicity, and four people with distant genetic distribution according to PCA of WGS data. The second round of quality control was performed on the specimen level, where we explored the snRNA-seq data quality of each specimen. Since some participants had replicated specimens, we also examined the experimental batch information of specimens to retain only one specimen for each sample that has the best quality and is from the most frequently used batches. Four parameters were calculated based on the snRNA-seq data for quality control, which are: 1) total number of cells in each specimen; 2) median of the total UMI counts in each specimen (median of nCount_RNA); 3) median of the expressed number of genes in each specimen (median of nFeature_RNA); 4) number of cells in each cell type/cluster. We excluded specimens if they satisfied any of these criteria: 1) total number of cells is less than 100; 2) median UMI count; 3) median expressed features are less than 1000; and 4) number of cells in any of the six cell types we study is less than 10 cells. If a participant has replicated specimens, we selected the one with a higher abundance in measurements and from a batch where most specimen samples were sequenced. As a result, one person with low-quality specimens in snRNA-seq was excluded, and all specimens from the MAP study were excluded since they had a lower quality cell type level. Participants who were diagnosed with AD were selected based on a neuropathological diagnosis using semiquantitative estimates of neuritic plaque density, as recommended by the Consortium to Establish a Registry for Alzheimer's Disease (CERAD). As a result, 272 AD patients with sufficient data quality remained for constructing gene regulatory networks. **Table 1** contains a summary of demographic information for the 272 participants retained after quality control.

#### S3. Single-nuclei Gene Expression Data Preparation

We performed cell type classification on all 272 participants using the same procedure as described in the ROSMAP study.[3] Briefly, the count matrix was normalized using SCTransform with the most variable 4000 genes. Dimension reduction was performed with RunPCA with npcs=50, and the cells were clustered using FindNeighbors (dims=1:50) and FindClusters (resolution=1.5, algorithm=4), and a UMAP plot was generated with RunUMAP (dims=1:50, min.dist=0.1). The cell clusters went through both automatic annotation, using a weighted elastic-net logistic regression with 24 donors from a previous DLPFC atlas as the reference database, and functional annotation to assign cells to major brain cell types.[3–5] Finally, we obtained seven major cell types, including excitatory neurons (375,092 nuclei), inhibitory neurons (148,224 nuclei), astrocytes (131,008 nuclei), microglia (49,263 nuclei), oligodendrocytes (202,714 nuclei), oligodendrocyte precursor cells (OPCs; 34,936 nuclei), and endothelial cells (6,031 nuclei). (**Figure 2A**) Endothelial cells were detected in 271 samples, with an average of only 22 nuclei available per individual, and therefore were excluded from the network construction.

#### S4. Aggregation of Single-nucleus RNA-seq and Covariates Adjustment

To construct cell-type-specific GRNs using both snRNA-seq and genotype, the sparse snRNA-seq data need to be aggregated to generate individual-level gene expression for each cell type. The aggregated counts were calculated by summing the expression level of all cells assigned to the cell type for each individual. Confounding covariates were adjusted using a generalized linear mixed model (GLMM) with a Poisson distribution on the aggregated counts to remove any unwanted variations. Age, gender, post-mortem interval (PMI), study cohort (ROS or MAP), and APOE genotype were used as fixed effects, while the batch effects of specimens were included in the model as random effects. The number of nuclei in each individual was log-transformed and was used as an offset term to account for variations in cellular sampling depth. The model could be formulated as follows:

$$\log(Y_{ij}) = \beta_0 + \beta_1 Age_i + \beta_2 Gender_i + \beta_3 PMI_i + \beta_4 Study_i + \beta_5 APOE_i + b_j + \log(N_i),$$

where $Y_{ij}$ is the aggregated gene expression counts in individual $i$ and batch $j$, $\beta$s are fixed effects coefficients, $b_j$ represents the random effect of batch $j$ and $\log(N_i)$ is the offset term where $N_i$ is the number of nuclei in each individual. After fitting this model, we extracted the working residuals as the normalized gene expression for the subsequent analyses.

### S5. Genotype Preprocessing and Imputation

The construction of GRN with genetic markers as instrumental variables requires careful preprocessing of genotype data to ensure data quality. We excluded SNPs with multiallelic or duplicated alleles, keeping only the most reliable biallelic SNPs for analysis. These remaining SNPs went through strict filtering criteria, such that we retain only SNPs with a missing rate < 0.1, passed the Hardy-Weinberg equilibrium test with a p-value cutoff of 1E-6, and with a minimum minor allele count greater than five across all individuals. At the sample level, we also examined all individuals with a genotype missing rate greater than 0.2 to ensure enough genomic coverage. All quality control steps were performed using PLINK (v1.9).[6]

After the above quality control, we imputed the remaining missing genotype using IMPUTE2 software with 1000 Genome Phase 3 as the reference panel.[7] To efficiently run computationally intensive jobs on the whole genome, we generated parallel scripts by dividing the genome into windows of 100kb each, which were simultaneously submitted to high-performance computational servers to reduce the computational time. After the imputation jobs were completed, they were recombined by windows into the whole genome data.

The imputed genotype was further filtered with an imputation quality score of 0.5, and we then repeated the above quality control. The final genotype, which has passed these two rounds of quality control and imputation, was prepared for the next step of GRN construction.

### S6. Construction of Gene Regulatory Networks

We constructed cell-type-specific gene regulatory networks via causal inference using SIGNET. This method incorporated genetic markers as instrumental variables into a transcriptome-wide Mendelian Randomization framework to infer causal gene-gene regulations, which outperformed commonly used correlation-based approaches in revealing causation instead of correlation.

The SIGNET pipeline contains two methodological steps. The first step aims to identify appropriate instrumental variables (IVs) for each target gene. We scanned for variants located within the cis-regulatory region of each gene, which is defined to be 1000 base pairs upstream and downstream of the gene's genomic region. These variants were stratified by their minor allele frequency (MAF) levels into three categories: common variants (MAF≥0.05), low-frequency variants (0.01≤MAF<0.05), and rare variants (MAF<0.01). Common variants were directly tested using a univariate regression model to select IVs. In contrast, low and rare variants were aggregated into 50 and 100 variants per window sizes, respectively, and went through an adaptive burden test, aSum test, to evaluate their role as potential IVs.

In the second step, we performed a transcriptome-wide multiple Mendelian Randomization on each cell type to infer causal regulatory relationships given the genomic IVs. This was accomplished through a two-stage penalized least squares (2SPLS) approach. In Stage 1, we used the identified IVs to predict the gene expression with ridge regression, where the tuning parameter was optimized using generalized cross-validation (GCV). In Stage 2, we applied adaptive LASSO regression to identify significant causal regulations between genes. This step was carried out with parallel computing across genes at each stage to improve computational efficiency.

To ensure the liability of the identified regulations, we bootstrapped 1000 datasets and constructed a GRN for each dataset. Only regulatory edges appeared more than 95% of the GRNs were retained for further investigation.

Upon completion of the above approaches, we obtained a sparse adjacent matrix representing the GRN of each cell type, with each entry (i,j) indicating the regulatory effect of gene i on gene j, as well as a table with the bootstrap frequency for each regulation. Due to the extensive size and complexity of the full network, we further decomposed it into sub-modules based on the network connectivity using a community detection algorithm.[8] For subsequent analyses of AD-related regulatory mechanisms, we focused on the largest five sub-networks with the most densely connected regulations, which are likely the core modules to represent the cellular functions.

### S7. Identification of Hub Genes from GRNs

Following [9], we calculate the coefficient of determination to quantify the proportion of variations explained by its regulators for each $i$ , that is, when properly standardized

$$\boldsymbol{R}_i^2 = \mathbf{1} - \left\|\boldsymbol{Y}_i - \sum_{j\neq i}\boldsymbol{Y}_j\hat{\boldsymbol{\gamma}}_{ij}\right\|_2^2 / \|\boldsymbol{Y}_i\|_2^2 \text{ ,}$$

In addition, we define the edgewise variation of gene $i$ explains its target $j$, as

$$C_{ij}^2 = \mathbf{1} - \left\|\boldsymbol{Y}_j - \boldsymbol{Y}_i\,\hat{\boldsymbol{\gamma}}_{ji}\right\|_2^2 / \left\|\boldsymbol{Y}_j\right\|_2^2$$

calculate the sum of the proportion of variations of its targets for each gene $i$ , where we have

$$C_i^2 = \sum_{j=1}^{p} C_{ij}^2.$$

where $Y_i$ encodes the gene expression vector for gene $i$ and $\hat{\gamma}_{ij}$ represents the estimated causal effect of the $j$-th gene on the $i$-th gene.

### S8. Pathway Enrichment Analysis Across Cell Types

To understand the functional implications of the constructed cell-type-specific GRNs, we performed enrichment analysis using the GO and KEGG pathways and identified distinct pathways and regulatory patterns across cell types. (**Figure S5A**) Excitatory and inhibitory neurons showed enrichment in similar GO and KEGG terms, especially in neuron synaptic conjunctions. However, inhibitory neurons are enriched in a few additional monoatomic ion channel activities in GO molecular functions than excitatory neurons. In terms of enrichment significance, inhibitory neurons had higher enrichment p-values for GO terms than excitatory neurons. In contrast, excitatory neurons are extremely more significant in KEGG pathways, especially for pathways related to human neurodegenerative diseases such as AD, Parkinson's disease, and Huntington's disease.

For glial cells, astrocytes display enrichment in similar pathways compared to neurons, especially in GO pathways related to biological processes and cellular components, while their enrichment pattern in molecular function and KEGG pathways is more diverse. More specifically, astrocytes are the only cell type that was highly enriched in regulatory activities of small GTPase mediated signal transduction in GO terms. Being the smallest cell population with the sparsest network, OPCs were mainly enriched in cellular components of GO pathways, with the only enriched pathway in KEGG being cell adhesive molecules. In contrast, none of the top pathways in oligodendrocytes were associated with cellular components, which suggests the difference in the roles of OPCs compared with oligodendrocytes. Despite microglia having a larger network size than OPCs, it was enriched in the smallest number of pathways overall. Microglia were mainly enriched in Th17 and myeloid cell differentiation and GTPase regulator activities, which are key pathways related to neuroinflammation or immune function.

Several enriched pathways were identified across multiple cell types, while the regulatory structures surrounding the common and hub genes present different patterns. This may suggest their cell-type-specific roles in the shared biological functions related to AD. As an example, alcoholic/steroid/cholesterol metabolic pathways were identified in three cell types: astrocytes, inhibitory neurons, and OPCs. (**Figure S3A**) Gene *MSMO1* acted as a driving hub that mainly regulated other genes in both inhibitory neurons and OPCs; however, in astrocytes, *MSMO1* only acted as a target that was regulated by upstream genes. Another shared pathway with similar interactions is the protein folding pathway, which was enriched in microglia, oligodendrocytes, and OPCs. (**Figure S5B**) Gene *HSPH1* appeared as the sole driving hub in microglia and regulated the remaining nine genes in this pathway. On the contrary, in oligodendrocytes and OPCs, *HSPH1* didn't play an important role in the networks and mainly reacted to other genes in the heat shock protein (HSP) family. Besides, the connections related to protein folding are more complex in oligodendrocytes with a larger cluster of genes, compared to microglia and OPCs. (**Figure S5C**)

### S9. IPA Analysis

We employed IPA (Ingenuity Pathway Analysis) to construct gene regulatory networks based on its curated database using our inferred genes from each of the top subnetworks. We incorporated both direct and indirect relationships. For each subnetwork, we included up to 70 genes for each subnetwork specifically on relationships from the human nervous system for Tissue & Cancer Lines, utilizing IPA's extensive curated database to contextualize and validate our findings.

### Supplementary Tables and Figures

#### [Supplementary Tables]

#### Table S1. Summary of regulations in GRNs and sub-networks of each cell type

This table provides an overview of the complexity of GRNs in each cell type, split by sub-networks. Each entry indicates the number of regulations in the row categories of the corresponding cell type, followed by the number of genes involved in these regulations within parentheses. The five largest sub-networks accounted for 50.93-65.89% of the full networks, while the largest network accounted for 27.10-61.96% of the full networks. The largest sub-network of microglia shows the lowest percentage of 27.10% followed by the largest sub-network of OPCs with 38.88%.

| # of Regulations (# of Genes) | Neurons | | Glial Cells | | | |
|---|---|---|---|---|---|---|
| | Exc. | Inh. | Ast. | Oli. | OPC. | Mic. |
| **Full network** | 5910 (5756) | 2428 (2522) | 2446 (2437) | 1290 (1323) | 697 (732) | 871 (865) |
| **Sum of the top 5 sub-networks** | 3894 (3049) | 1476 (1158) | 1547 (1207) | 778 (596) | 355 (273) | 444 (340) |
| **The largest sub-network** | 3662 (2848) | 1380 (1081) | 1390 (1079) | 725 (543) | 271 (204) | 236 (175) |
| **The 2nd largest sub-network** | 164 (139) | 32 (28) | 75 (58) | 25 (23) | 24 (20) | 96 (73) |
| **The 3rd largest sub-network** | 27 (25) | 29 (19) | 31 (25) | 12 (12) | 23 (19) | 53 (45) |
| **The 4th largest sub-network** | 25 (21) | 17 (15) | 28 (23) | 10 (11) | 19 (18) | 35 (28) |
| **The 5th largest sub-network** | 16 (16) | 18 (15) | 23 (22) | 6 (7) | 18 (12) | 24 (19) |

**Table S2. All genes in the networks of all cell types (See Excel table)**

This Excel table contains a detailed list of all nodes included in the full networks for all six cell types. Each row is a gene. This table included logical variables indicating whether a gene is categorized as a hub gene, regulatory hub (outhub), or responsive hub (inhub) under a 0.95 quantile cutoff threshold. It also reported the exact values for degree (both indegree and outdegree), $R^2$ and $C^2$ values.

**Table S3. Comparison of hub genes across cell types (Excel table)**

The lists of hub genes in all cell types were compared to find overlapping and cell-type unique hubs. There were no hubs shared across all cell types, and no hubs were found among the glial cells. Only neuron cells, excitatory and inhibitory neurons, exhibited a list of shared hubs. The cell-type unique hubs were defined to be genes that appeared solely in the target cell type, and not in any other cell types. There were separate tables for each of the following lists: common hubs between neurons, cell-type unique hubs for each of the six cell types. The table in this file has the same format as **Table S2**.

**Table S4. Enriched GO and KEGG pathways of all hub genes and their targets in excitatory neurons, corresponding to the IDs in Figure 4B.**

This table reports the enriched GO and KEGG pathways of all hub genes and their nearest two neighboring genes in excitatory neurons, from the direction of upstream regulators to their targets. Source pathways are the significantly enriched pathways of upstream regulators, and target pathways are the significantly enriched pathways of downstream targets. Degree indicates the number of gene regulations involved in each pair of pathways. The table was sorted by degree in descending order. A heatmap illustration of this table can be found in **Figure 4B**.

A. GO pathways

| Source Pathways | Target Pathways | Source Pathway Names | Target Pathway Names | Degree |
|---|---|---|---|---|
| GO:1902495 | GO:0005743 | GO:transmembrane transporter complex | GO:mitochondrial inner membrane | 39 |
| GO:1990351 | GO:0005743 | GO:transporter complex | GO:mitochondrial inner membrane | 39 |
| GO:0097060 | GO:0097060 | GO:synaptic membrane | GO:synaptic membrane | 35 |
| GO:0042391 | GO:0097060 | GO:regulation of membrane potential | GO:synaptic membrane | 32 |
| GO:0097060 | GO:0043025 | GO:synaptic membrane | GO:neuronal cell body | 31 |
| GO:0097060 | GO:0050808 | GO:synaptic membrane | GO:synapse organization | 30 |
| GO:0097060 | GO:0099572 | GO:synaptic membrane | GO:postsynaptic specialization | 30 |
| GO:0043025 | GO:0043025 | GO:neuronal cell body | GO:neuronal cell body | 29 |
| GO:0098978 | GO:0043025 | GO:glutamatergic synapse | GO:neuronal cell body | 29 |
| GO:0050804 | GO:0097060 | GO:modulation of chemical synaptic transmission | GO:synaptic membrane | 29 |
| GO:0099177 | GO:0097060 | GO:regulation of trans-synaptic signaling | GO:synaptic membrane | 29 |
| GO:0043025 | GO:0098978 | GO:neuronal cell body | GO:glutamatergic synapse | 29 |
| GO:0097060 | GO:0098984 | GO:synaptic membrane | GO:neuron to neuron synapse | 29 |
| GO:0050804 | GO:0050804 | GO:modulation of chemical synaptic transmission | GO:modulation of chemical synaptic transmission | 28 |
| GO:0097060 | GO:0050804 | GO:synaptic membrane | GO:modulation of chemical synaptic transmission | 28 |
| GO:0099177 | GO:0050804 | GO:regulation of trans-synaptic signaling | GO:modulation of chemical synaptic transmission | 28 |
| GO:0043025 | GO:0097060 | GO:neuronal cell body | GO:synaptic membrane | 28 |
| GO:0050808 | GO:0097060 | GO:synapse organization | GO:synaptic membrane | 28 |
| GO:0050804 | GO:0099177 | GO:modulation of chemical synaptic transmission | GO:regulation of trans-synaptic signaling | 28 |
| GO:0097060 | GO:0099177 | GO:synaptic membrane | GO:regulation of trans-synaptic signaling | 28 |
| GO:0099177 | GO:0099177 | GO:regulation of trans-synaptic signaling | GO:regulation of trans-synaptic signaling | 28 |

| | | | | |
|---|---|---|---|---|
| GO:0034329 | GO:0050808 | GO:cell junction assembly | GO:synapse organization | 27 |
| GO:0034329 | GO:0097060 | GO:cell junction assembly | GO:synaptic membrane | 27 |
| GO:0099572 | GO:0097060 | GO:postsynaptic specialization | GO:synaptic membrane | 27 |
| GO:0097060 | GO:0014069 | GO:synaptic membrane | GO:postsynaptic density | 26 |
| GO:0097060 | GO:0032279 | GO:synaptic membrane | GO:asymmetric synapse | 26 |
| GO:0050804 | GO:0050808 | GO:modulation of chemical synaptic transmission | GO:synapse organization | 26 |
| GO:0099177 | GO:0050808 | GO:regulation of trans-synaptic signaling | GO:synapse organization | 26 |
| GO:0042391 | GO:0098984 | GO:regulation of membrane potential | GO:neuron to neuron synapse | 26 |
| GO:0042391 | GO:0043025 | GO:regulation of membrane potential | GO:neuronal cell body | 25 |

### B. KEGG pathways

| Source Pathways | Target Pathways | Source Pathway Names | Target Pathway Names | Degree |
|---|---|---|---|---|
| hsa05022 | hsa05022 | KEGG:Pathways of neurodegeneration - multiple diseases | KEGG:Pathways of neurodegeneration - multiple diseases | 72 |
| hsa05010 | hsa05022 | KEGG:Alzheimer disease | KEGG:Pathways of neurodegeneration - multiple diseases | 69 |
| hsa05020 | hsa05022 | KEGG:Prion disease | KEGG:Pathways of neurodegeneration - multiple diseases | 69 |
| hsa05022 | hsa05010 | KEGG:Pathways of neurodegeneration - multiple diseases | KEGG:Alzheimer disease | 68 |
| hsa05012 | hsa05022 | KEGG:Parkinson disease | KEGG:Pathways of neurodegeneration - multiple diseases | 68 |
| hsa05012 | hsa05012 | KEGG:Parkinson disease | KEGG:Parkinson disease | 66 |
| hsa05022 | hsa05012 | KEGG:Pathways of neurodegeneration - multiple diseases | KEGG:Parkinson disease | 66 |
| hsa05022 | hsa05014 | KEGG:Pathways of neurodegeneration - multiple diseases | KEGG:Amyotrophic lateral sclerosis | 66 |
| hsa05014 | hsa05022 | KEGG:Amyotrophic lateral sclerosis | KEGG:Pathways of neurodegeneration - multiple diseases | 66 |
| hsa05010 | hsa05010 | KEGG:Alzheimer disease | KEGG:Alzheimer disease | 65 |
| hsa05020 | hsa05014 | KEGG:Prion disease | KEGG:Amyotrophic lateral sclerosis | 65 |
| hsa05012 | hsa05010 | KEGG:Parkinson disease | KEGG:Alzheimer disease | 64 |
| hsa05020 | hsa05010 | KEGG:Prion disease | KEGG:Alzheimer disease | 64 |
| hsa05020 | hsa05012 | KEGG:Prion disease | KEGG:Parkinson disease | 64 |
| hsa05010 | hsa05014 | KEGG:Alzheimer disease | KEGG:Amyotrophic lateral sclerosis | 64 |
| hsa05012 | hsa05014 | KEGG:Parkinson disease | KEGG:Amyotrophic lateral sclerosis | 64 |
| hsa05014 | hsa05014 | KEGG:Amyotrophic lateral sclerosis | KEGG:Amyotrophic lateral sclerosis | 64 |
| hsa05012 | hsa05016 | KEGG:Parkinson disease | KEGG:Huntington disease | 64 |
| hsa05022 | hsa05016 | KEGG:Pathways of neurodegeneration - multiple diseases | KEGG:Huntington disease | 64 |
| hsa05022 | hsa05020 | KEGG:Pathways of neurodegeneration - multiple diseases | KEGG:Prion disease | 64 |
| hsa05014 | hsa05010 | KEGG:Amyotrophic lateral sclerosis | KEGG:Alzheimer disease | 63 |
| hsa05010 | hsa05012 | KEGG:Alzheimer disease | KEGG:Parkinson disease | 63 |
| hsa05014 | hsa05012 | KEGG:Amyotrophic lateral sclerosis | KEGG:Parkinson disease | 63 |
| hsa05020 | hsa05020 | KEGG:Prion disease | KEGG:Prion disease | 63 |
| hsa05010 | hsa05016 | KEGG:Alzheimer disease | KEGG:Huntington disease | 62 |
| hsa05014 | hsa05016 | KEGG:Amyotrophic lateral sclerosis | KEGG:Huntington disease | 62 |
| hsa05010 | hsa05020 | KEGG:Alzheimer disease | KEGG:Prion disease | 62 |

| | | | | |
|---|---|---|---|---|
| hsa05012 | hsa05020 | KEGG:Parkinson disease | KEGG:Prion disease | 62 |
| hsa05020 | hsa05016 | KEGG:Prion disease | KEGG:Huntington disease | 61 |
| hsa05014 | hsa05020 | KEGG:Amyotrophic lateral sclerosis | KEGG:Prion disease | 61 |

**Table S5. Enriched GO and KEGG pathways of TF hubs and their targets in excitatory neurons, corresponding to the IDs in Figure S4.**

This table reports the enriched GO and KEGG pathways of TF hub genes and their nearest two neighboring genes in excitatory neurons, from the direction of upstream regulators to their targets. Source pathways are the significantly enriched pathways of upstream regulators, and target pathways are the significantly enriched pathways of downstream targets. Degree indicates the number of gene regulations involved in each pair of pathways. The table was sorted by degree in descending order. A heatmap illustration of this table can be found in **Figure S4**.

### A. GO pathway names

| Source Pathways | Target Pathways | Source Pathway Names | Target Pathway Names | Degree |
|---|---|---|---|---|
| GO:0050808 | GO:0003012 | GO:synapse organization | GO:muscle system process | 9 |
| GO:0010563 | GO:0043409 | GO:negative regulation of phosphorus metabolic process | GO:negative regulation of MAPK cascade | 7 |
| GO:0042326 | GO:0043409 | GO:negative regulation of phosphorylation | GO:negative regulation of MAPK cascade | 7 |
| GO:0045936 | GO:0043409 | GO:negative regulation of phosphate metabolic process | GO:negative regulation of MAPK cascade | 7 |
| GO:0001933 | GO:0050804 | GO:negative regulation of protein phosphorylation | GO:modulation of chemical synaptic transmission | 7 |
| GO:0010563 | GO:0050804 | GO:negative regulation of phosphorus metabolic process | GO:modulation of chemical synaptic transmission | 7 |
| GO:0042326 | GO:0050804 | GO:negative regulation of phosphorylation | GO:modulation of chemical synaptic transmission | 7 |
| GO:0045936 | GO:0050804 | GO:negative regulation of phosphate metabolic process | GO:modulation of chemical synaptic transmission | 7 |
| GO:0001933 | GO:0050808 | GO:negative regulation of protein phosphorylation | GO:synapse organization | 7 |
| GO:0010563 | GO:0050808 | GO:negative regulation of phosphorus metabolic process | GO:synapse organization | 7 |
| GO:0042326 | GO:0050808 | GO:negative regulation of phosphorylation | GO:synapse organization | 7 |
| GO:0045936 | GO:0050808 | GO:negative regulation of phosphate metabolic process | GO:synapse organization | 7 |
| GO:0034702 | GO:0097060 | GO:monoatomic ion channel complex | GO:synaptic membrane | 7 |
| GO:1902495 | GO:0097060 | GO:transmembrane transporter complex | GO:synaptic membrane | 7 |
| GO:1990351 | GO:0097060 | GO:transporter complex | GO:synaptic membrane | 7 |
| GO:0001933 | GO:0099177 | GO:negative regulation of protein phosphorylation | GO:regulation of trans-synaptic signaling | 7 |
| GO:0010563 | GO:0099177 | GO:negative regulation of phosphorus metabolic process | GO:regulation of trans-synaptic signaling | 7 |

| | | | | |
|---|---|---|---|---|
| GO:0042326 | GO:0099177 | GO:negative regulation of phosphorylation | GO:regulation of trans-synaptic signaling | 7 |
| GO:0045936 | GO:0099177 | GO:negative regulation of phosphate metabolic process | GO:regulation of trans-synaptic signaling | 7 |
| GO:0014069 | GO:0003012 | GO:postsynaptic density | GO:muscle system process | 6 |
| GO:0032279 | GO:0003012 | GO:asymmetric synapse | GO:muscle system process | 6 |
| GO:0098984 | GO:0003012 | GO:neuron to neuron synapse | GO:muscle system process | 6 |
| GO:0099572 | GO:0003012 | GO:postsynaptic specialization | GO:muscle system process | 6 |
| GO:0001933 | GO:0007611 | GO:negative regulation of protein phosphorylation | GO:learning or memory | 6 |
| GO:0010563 | GO:0007611 | GO:negative regulation of phosphorus metabolic process | GO:learning or memory | 6 |
| GO:0042326 | GO:0007611 | GO:negative regulation of phosphorylation | GO:learning or memory | 6 |
| GO:0045936 | GO:0007611 | GO:negative regulation of phosphate metabolic process | GO:learning or memory | 6 |
| GO:0001933 | GO:0043025 | GO:negative regulation of protein phosphorylation | GO:neuronal cell body | 6 |
| GO:0006813 | GO:0043025 | GO:potassium ion transport | GO:neuronal cell body | 6 |
| GO:0010563 | GO:0043025 | GO:negative regulation of phosphorus metabolic process | GO:neuronal cell body | 6 |

### B. KEGG pathway names

| Source Pathways | Target Pathways | Source Pathway Names | Target Pathway Names | Degree |
|---|---|---|---|---|
| hsa05031 | hsa00534 | KEGG:Amphetamine addiction | KEGG:Glycosaminoglycan biosynthesis - heparan sulfate / heparin | 3 |
| hsa04082 | hsa00534 | KEGG:Neuroactive ligand signaling | KEGG:Glycosaminoglycan biosynthesis - heparan sulfate / heparin | 2 |
| hsa04713 | hsa00534 | KEGG:Circadian entrainment | KEGG:Glycosaminoglycan biosynthesis - heparan sulfate / heparin | 2 |
| hsa04724 | hsa00534 | KEGG:Glutamatergic synapse | KEGG:Glycosaminoglycan biosynthesis - heparan sulfate / heparin | 2 |
| hsa04728 | hsa00534 | KEGG:Dopaminergic synapse | KEGG:Glycosaminoglycan biosynthesis - heparan sulfate / heparin | 2 |
| hsa04261 | hsa04724 | KEGG:Adrenergic signaling in cardiomyocytes | KEGG:Glutamatergic synapse | 2 |

### Supplementary Figures

#### Figure S1. IPA network figures for each sub-network

Genes from each sub-network were submitted to IPA software to find overlaps between our GRNs and past studies in the IPA database. Genes identified in common were shown in these IPA figures, with connections indicating the identified relationship from past studies.

A. An overlapping network between IPA and our largest network of excitatory neurons.
B. An overlapping network between IPA and our largest network of inhibitory neurons.

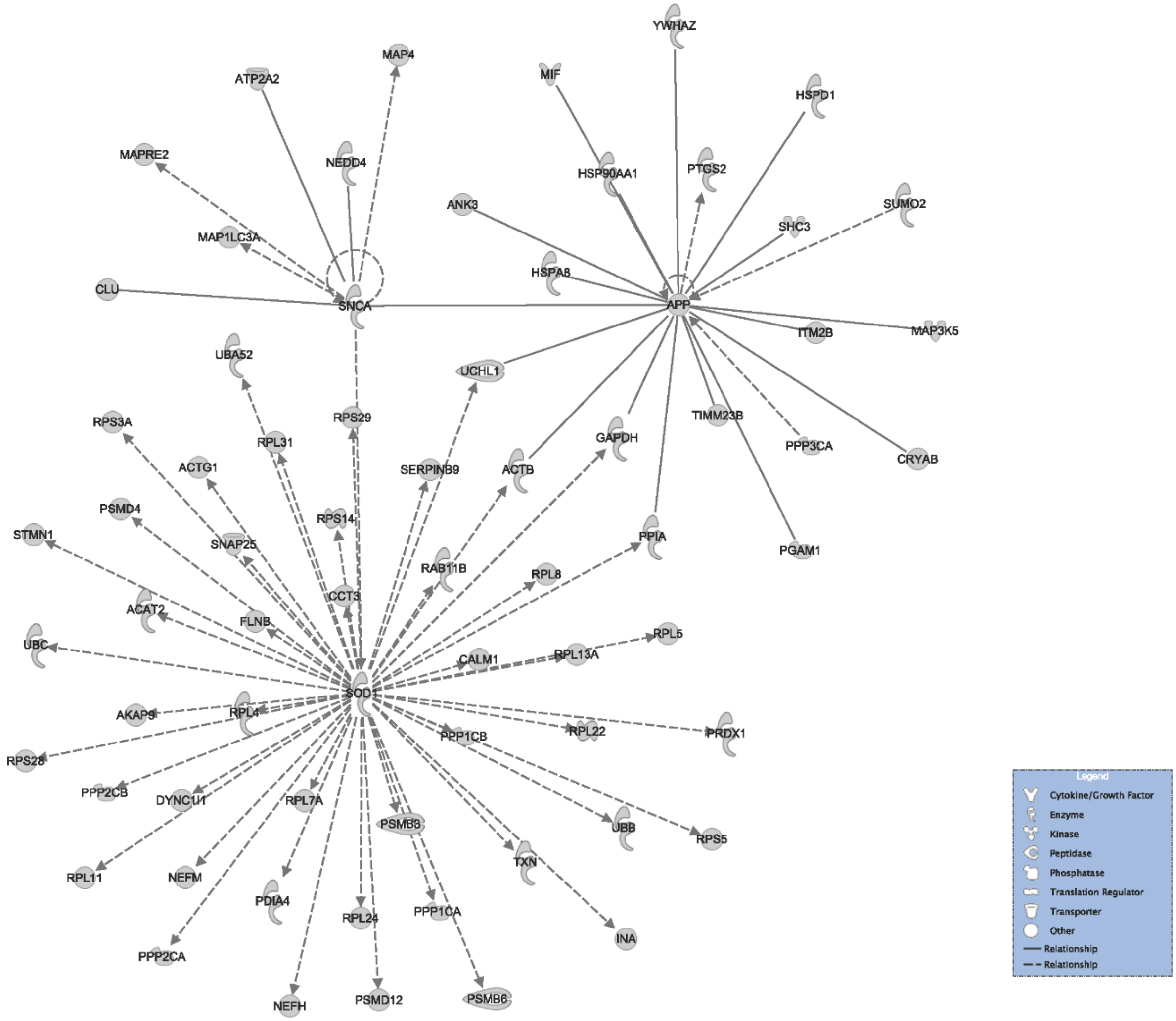

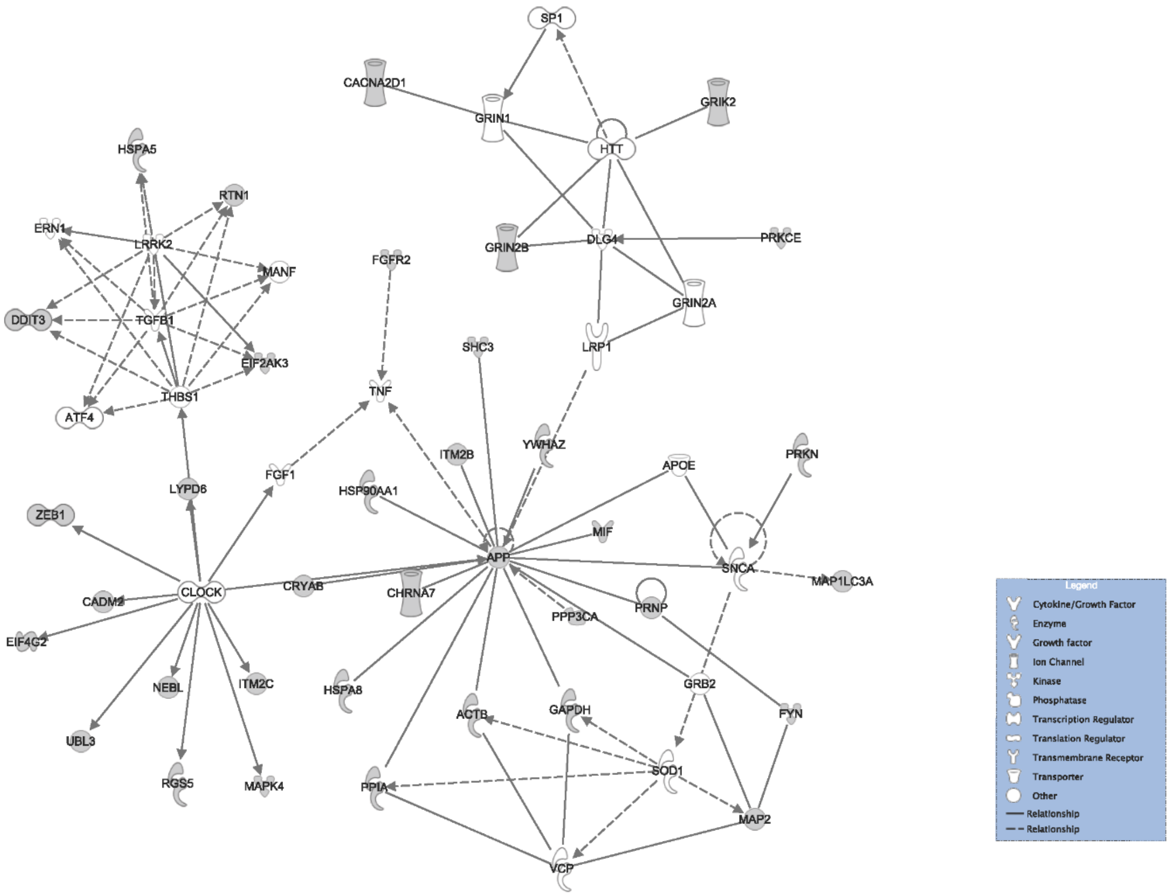

SP1
CACNA2D1
GRIN1
HTT
GRIK2
HSPA5
RTN1
ERN1
LRRK2
MANF
DDIT3
TGFB1
EIF2AK3
THBS1
ATF4
FGFR2
TNF
GRIN2B
DLG4
PRKCE
GRIN2A
SHC3
LRP1
ITM2B
YWHAZ
APOE
PRKN
FGF1
LYPD6
HSP90AA1
MIF
ZEB1
APP
SNCA
MAP1LC3A
CLOCK
CRYAB
CHRNA7
CADM2
PPP3CA
PRNP
EIF4G2
NEBL
ITM2C
HSPA8
GRB2
ACTB
GAPDH
FYN
UBL3
RGS5
MAPK4
PPIA
SOD1
MAP2
VCP
Legend
Cytokine/Growth Factor
Enzyme
Growth factor
Ion Channel
Kinase
Phosphatase
Transcription Regulator
Translation Regulator
Transmembrane Receptor
Transporter
Other
Relationship
Relationship

**Figure S2. UMAP plot of cell classification in the independent cohort**

A UMAP plot showing the cell populations in the UCI Multiomics Cohort. Each dot represents a single cell, and the cell clusters were annotated based on the canonical differential genes in the original study.[10] Pericyte/endothelia (PER.END) cells (in pink) were excluded from the analysis due to the restricted cell numbers and sample size. (Abbreviations: ASC: astrocytes; EX: excitatory neurons; INH: inhibitory neurons; MG: microglia; ODC: oligodendrocytes; OPC: oligodendrocyte progenitor cells; PER.END: pericyte/endothelia cells)

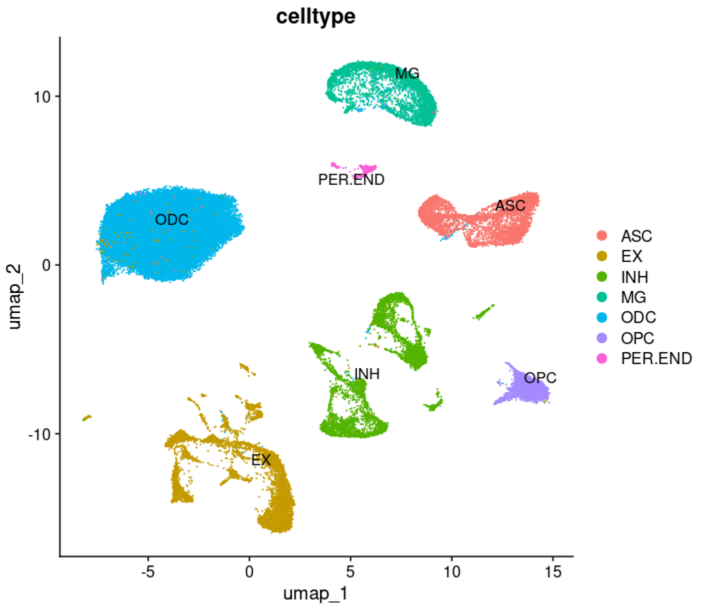

### Figure S3. Enriched GO and KEGG pathways for source and target genes, respectively

This tree figure shows the enriched GO and KEGG pathways from separate enrichment analysis for each source gene and target gene group. The top 2 panels are for GO pathways, and the bottom two panels are for KEGG pathways. The top 30 enriched pathways with more than 10 regulations were selected for this figure. Pathways with similar names and keywords were grouped into clusters, and the keywords of each cluster were summarized on the right label through text mining. The circles before the pathway name were colored by adjusted p-values from enrichment tests, and the number of genes determined their sizes within each pathway.

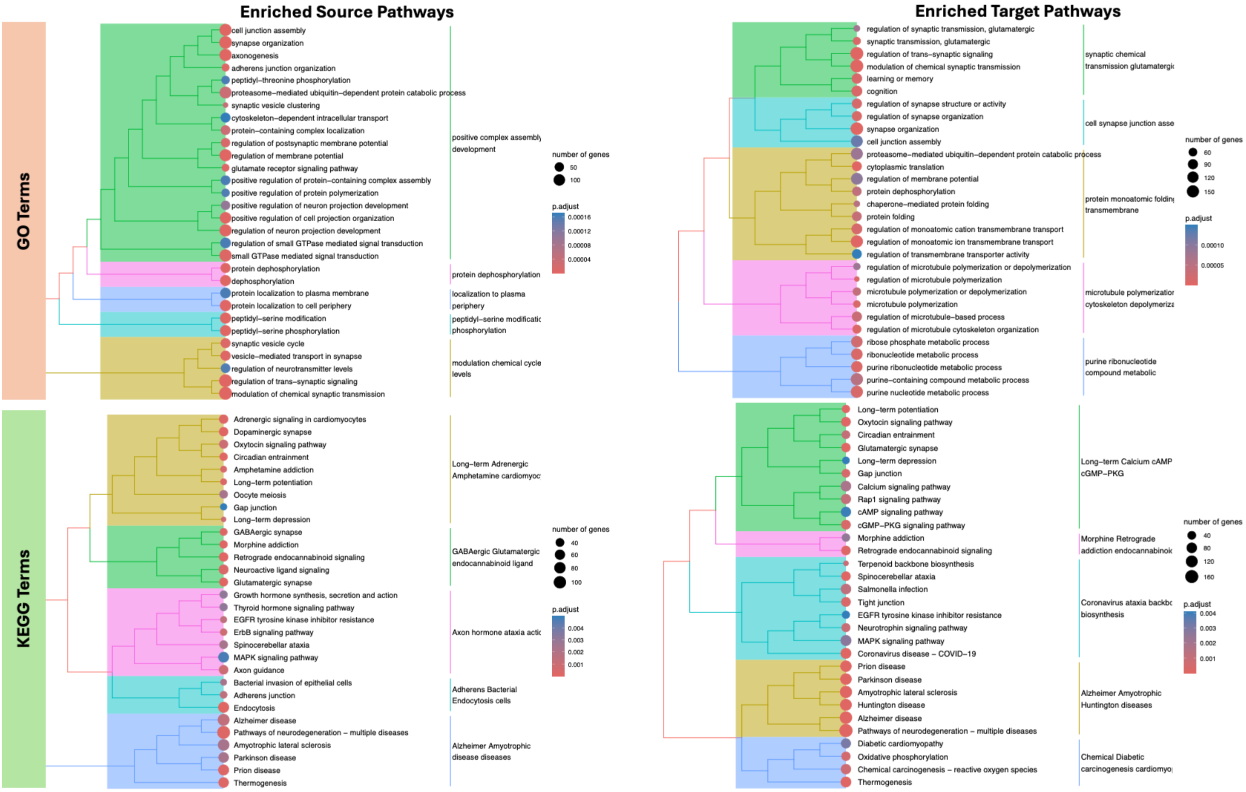

**Figure S4. Enriched GO/KEGG pathways of TF hubs and neighbors**

Heatmaps showing the pathway enrichment signals of TF hubs and their nearest two neighboring genes in excitatory neurons, from the direction of upstream regulators to their targets, for both GO and KEGG pathways. Only the top 30 significantly enriched pathways were included in the heatmap, with rows as enriched pathways of source genes and columns as enriched pathways of target genes. The blocks in the heatmap were colored based on the degree (i.e., the number of regulations) between a pair of pathways. A darker color indicates more regulations. Corresponding pathway names of the IDs can be found in **Table S5**.

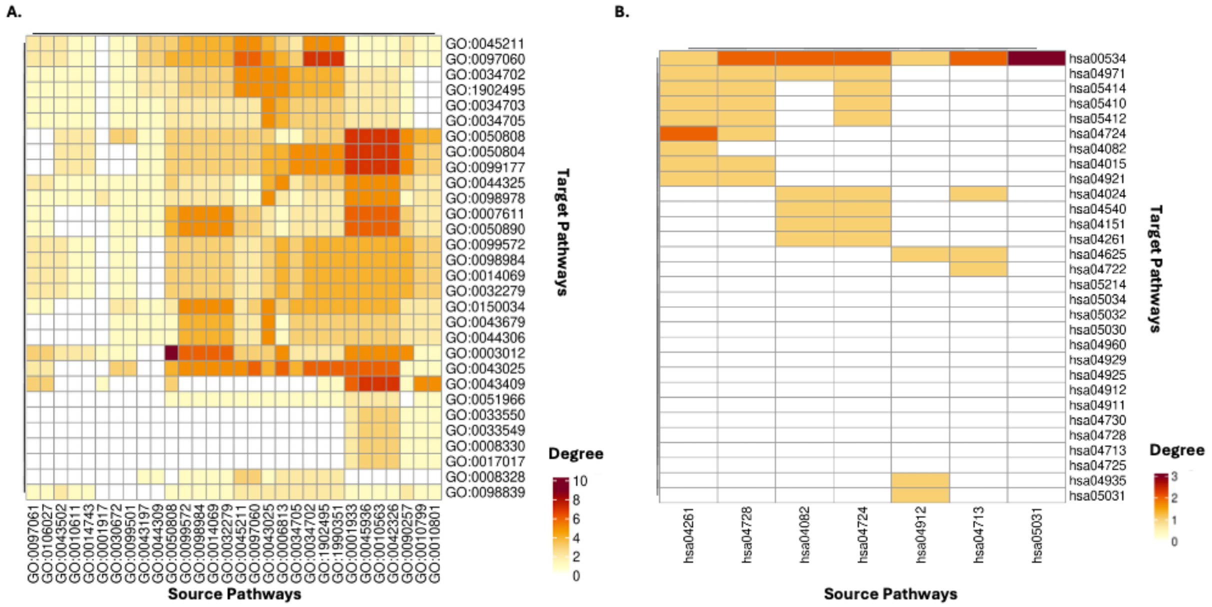

**Figure S5. Comparison of pathways and genes in the same pathway across cell types**

A. A heatmap showing the top enriched GO and KEGG pathways and their occurrence across cell types. The pathways shown in the figure were from the union of the top 5 most enriched pathways of all cell types by categories. The blocks were colored by the -log10 of the adjusted p-values. An empty block means the pathway was not enriched under a cutoff of adjusted p-values <10e-5 in the corresponding cell type.
B. Enriched genes and their regulations from the same pathway across different cell types-- alcoholic/steroid/cholesterol metabolic pathways
C. Enriched genes and their regulations from the same pathway across different cell types-- protein folding pathway

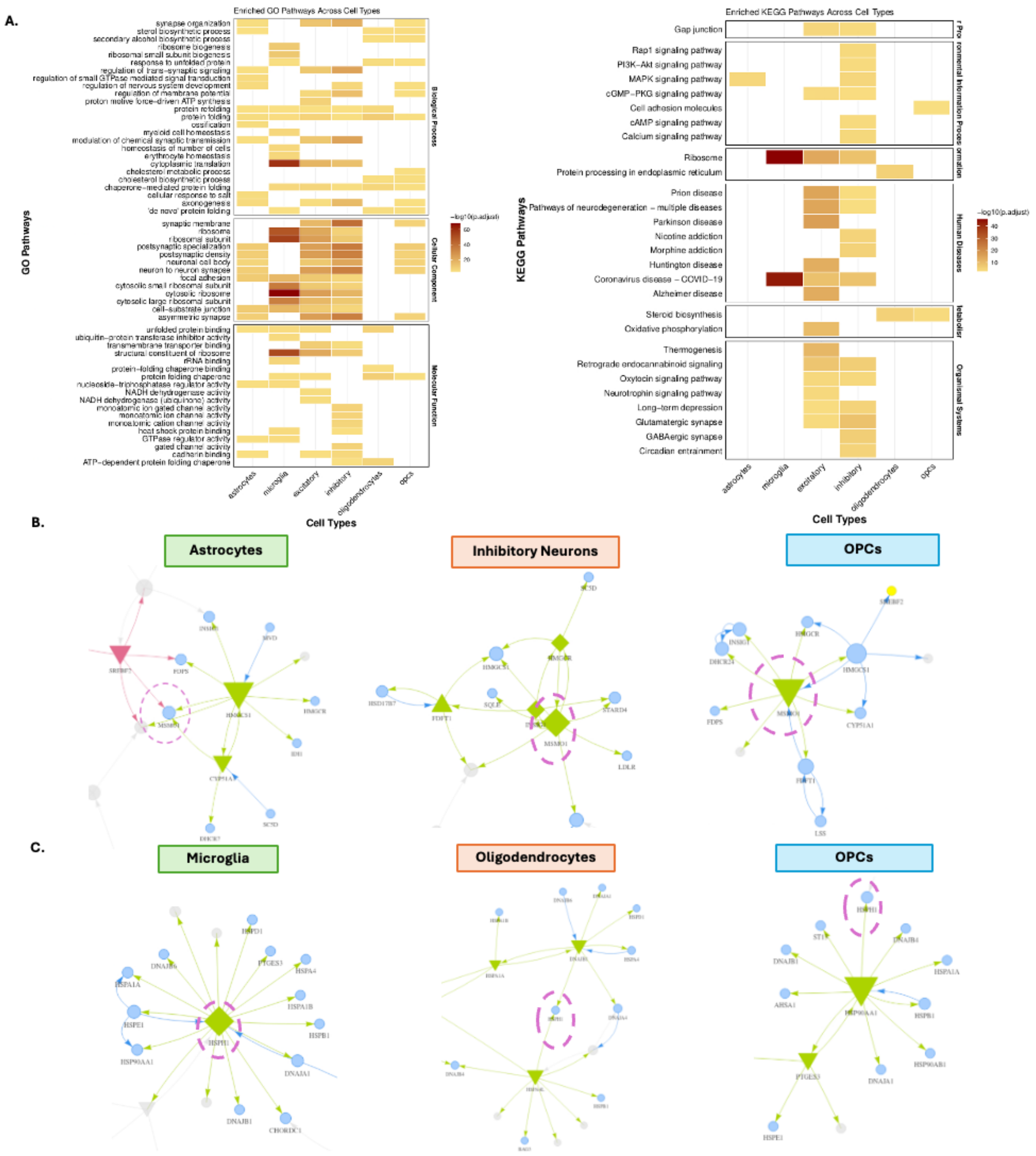

A.
Enriched GO Pathways Across Cell Types
GO Pathways
synapse organization
sterol biosynthetic process
secondary alcohol biosynthetic process
ribosome biogenesis
ribosomal small subunit biogenesis
response to unfolded protein
regulation of trans−synaptic signaling
regulation of small GTPase mediated signal transduction
regulation of nervous system development
regulation of membrane potential
proton motive force−driven ATP synthesis
protein refolding
protein folding
ossification
myeloid cell homeostasis
modulation of chemical synaptic transmission
homeostasis of number of cells
erythrocyte homeostasis
cytoplasmic translation
cholesterol metabolic process
cholesterol biosynthetic process
chaperone−mediated protein folding
cellular response to salt
axonogenesis
'de novo' protein folding
Biological Process
synaptic membrane
ribosome
ribosomal subunit
postsynaptic specialization
postsynaptic density
neuronal cell body
neuron to neuron synapse
focal adhesion
cytosolic small ribosomal subunit
cytosolic ribosome
cytosolic large ribosomal subunit
cell−substrate junction
asymmetric synapse
Cellular Component
unfolded protein binding
ubiquitin−protein transferase inhibitor activity
transmembrane transporter binding
structural constituent of ribosome
rRNA binding
protein−folding chaperone binding
protein folding chaperone
nucleoside−triphosphatase regulator activity
NADH dehydrogenase activity
NADH dehydrogenase (ubiquinone) activity
monoatomic ion gated channel activity
monoatomic ion channel activity
monoatomic cation channel activity
heat shock protein binding
GTPase regulator activity
gated channel activity
cadherin binding
ATP−dependent protein folding chaperone
Molecular Function
−log10(p.adjust)
60
40
20
astrocytes
microglia
excitatory
inhibitory
oligodendrocytes
opcs
Cell Types
Enriched KEGG Pathways Across Cell Types
KEGG Pathways
Gap junction
Rap1 signaling pathway
PI3K−Akt signaling pathway
MAPK signaling pathway
cGMP−PKG signaling pathway
Cell adhesion molecules
cAMP signaling pathway
Calcium signaling pathway
Ribosome
Protein processing in endoplasmic reticulum
Prion disease
Pathways of neurodegeneration − multiple diseases
Parkinson disease
Nicotine addiction
Morphine addiction
Huntington disease
Coronavirus disease − COVID−19
Alzheimer disease
Steroid biosynthesis
Oxidative phosphorylation
Thermogenesis
Retrograde endocannabinoid signaling
Oxytocin signaling pathway
Neurotrophin signaling pathway
Long−term depression
Glutamatergic synapse
GABAergic synapse
Circadian entrainment
Human Diseases
Organismal Systems
−log10(p.adjust)
40
30
20
10
astrocytes
microglia
excitatory
inhibitory
oligodendrocytes
opcs
Cell Types
B.
Astrocytes
Inhibitory Neurons
OPCs
C.
Microglia
Oligodendrocytes
OPCs
